# Loss of C3 and CD14 reduces region-specific neuroinflammation in a murine polytrauma model

**DOI:** 10.64898/2026.03.13.711583

**Authors:** Florian Olde Heuvel, Marica Pagliarini, Fan Sun, Ludmila Lupu, Zongren Zhao, Likun Cui, Rebecca Halbgebauer, Marco Mannes, Tobias Maria Boeckers, Egil Lien, Tom E. Mollnes, Markus Huber-Lang, Francesco Roselli

**Affiliations:** Dept. of Neurology-UIm University, Ulm, DE; Institute of Clinical and Experimental Trauma Immunology-Ulm University Medical Center, Ulm, DE; Institute of Anatomy and Cell Biology, Ulm University, Ulm, Germany; Division of Infectious Diseases and Immunology, Department of Medicine, University of Massachusetts Chan Medical School, Worcester, MA, US; Dept of Immunology, Oslo University Hospital-Oslo; Research Laboratory, Nordland Hospital Trust, Bodoe, Norway; German Center for Neurodegenerative Diseases (DZNE)-Ulm, DE

**Author notes:** co-corresponding authors, Corresponding author: Prof. Dr. Markus Huber-Lang, Institute of Clinical and Experimental Trauma Immunology, Ulm University Medical Center, Center for Biomedical Research (ZBF), Helmholtzstrasse 8/1-89081 Ulm DE, Prof. Dr. Dr. Francesco Roselli, Dept. of Neurology, Ulm University, Center for Biomedical Research (ZBF), Helmholtzstrasse 8/1-89081 Ulm DE. co-first authors (Florian Olde Heuvel, Marica Pagliarini, Fan Sun).

**Keywords:** Delirium, microglia, complement, TNF, cortex, striatum, polytrauma

## Abstract

**Background:** Traumatic brain injury (TBI) together with non-cerebral injuries characterizes the TBI-polytrauma (P-TBI) constellation, which is associated with acute neurological deterioration, delirium and unfavourable prognosis. It is hypothesized that systemic inflammatory mediators my enhances the focal, cerebral neuroimmune reaction with overall detrimental consequences, in particular in terms of acute microglial reactivity.

**Methods:** We explored the role of the Complement factor 3 (C3) and of the TLR-co receptor cluster of differentiation (CD14) in a murine polytrauma model that involves a mild TBI together with femur fracture, blunt thorax trauma and resuscitated haemorrhagic shock, making use of mice genetically lacking either C3, CD14 or both.

**Results:** We show that P-TBI results in a rapid (4h) and brain-wide induction of inflammatory cytokines, although with distinct profiles (*TNF* and *CCL2* having brain-wide involvement and *IL-1*β restricted to ipsilateral cortex and striatum). *TNF* and *CCL2* mRNA as well as protein synthesis were upregulated in microglia upon P-TBI in cortex, hippocampus and striatum which was fully abolished in the *C3^−/−^CD14^−/−^*animals. The analysis of single-KO animals revealed that induction of *TNF* and *CCL2* was prevented in animals lacking C3, but not CD14, in the contralateral cortex and striatum, with an abolishment in hippocampus in mice lacking both C3 and CD14. In the cortical area of focal lesion neither C3 nor CD14 affected the induction of pro-inflammatory cytokines.

**Conclusion:** Thus, C3 and CD14 are dispensable for the acute cytokine response to P-TBI in the site of injury but play differential roles across the cortex, hippocampus and striatum for the induction of cytokines in the non-injured parenchyma and in particular in microglia. Thus, interventions on C3 (mainly) and/or CD14 may reduce the encephalopathy risk associated with P-TBI but not the acute response in the injury site, where additional DAMP signalling may offer redundant activation pathways.

## Introduction

Polytrauma (PT), defined as the occurrence of injury in at least three body domains and several alterations in physiological parameters [1] is associated with multi-organ damage and high mortality [2]. The frequent involvement of the brain is mainly direct and focal, so that traumatic brain injury (TBI) is part of the initial PT constellation (PT including TBI, henceforth abbreviated as P-TBI). However, non-focal and rather related to secondary injury processes like altered level of consciousness, inattention and disorganised thinking as parts of the delirium diagnosis, and memory impairment are reported by more than half of P-TBI patients. These neurological consequences predict an overall worse long-term health status [3].

P-TBI is characterized by strong systemic immune activation initiated by the release of damage associated molecular patterns (DAMPs) which trigger signaling through a number of receptors, including Toll-like Receptors and their co-receptor CD14 [4]. TLR/CD14 signaling has been associated with secondary damage and organ dysfunction appearing in P-TBI patients and corresponding animal models [5,6].

The pathophysiology and thromboinflammatory response following P-TBI is also driven by biologically active products derived from multiple protease-dependent cascades, including the coagulation and complement system [7]. The latter is initiated within minutes after polytrauma and peaks within a few hours [8], thus constituting a major source of systemic inflammatory mediators in P-TBI and triggering further innate immunity processes [9].

Taken together, complement activation and TLR/CD14 signaling are considered prominent inducers of the systemic inflammatory response following P-TBI [10].

Posttraumatic neurologic complications including, encephalopathy and delirium are also frequently detected in other conditions characterized by systemic inflammation, such as sepsis [11] and major surgery [12]. In sepsis, the appearance of encephalopathy is linked to systemic complement activation [13] and associated with the induction of reactive microglia [14], and moreover with synaptic pruning and circuit dysfunction initiated by cerebral complement factors, such as C1q and C3 [15].

Therefore, we have hypothesized that microglial reactivity and central neuroimmune activation may be similarly occurring in the context of P-TBI. We set out to establish the early neuroinflammatory profile in P-TBI and its sensitivity to mice genetically lacking the complement factor C3 factor and the TLR CD14 co-signaling molecule.

## Results

### 1. Early cortical neuroinflammation in acute TBI/ polytrauma

We set out to explore the acute neuroinflammatory responses occurring in the cerebral cortex upon TBI associated with Polytrauma (P-TBI), in particular in a mouse model involving mild TBI, thorax trauma, bone fracture and haemorrhagic shock (HS) or sham (Fig. 1A). We focused on the 4h timepoint, compatible with timing for ICU admission of polytrauma patients [16]; at this stage, the overall architecture of the brain was preserved, and no necrotic/haemorrhagic lesions were observed in any of the brain samples on macroscopic examination (Fig. 1B).

**Figure 1:**
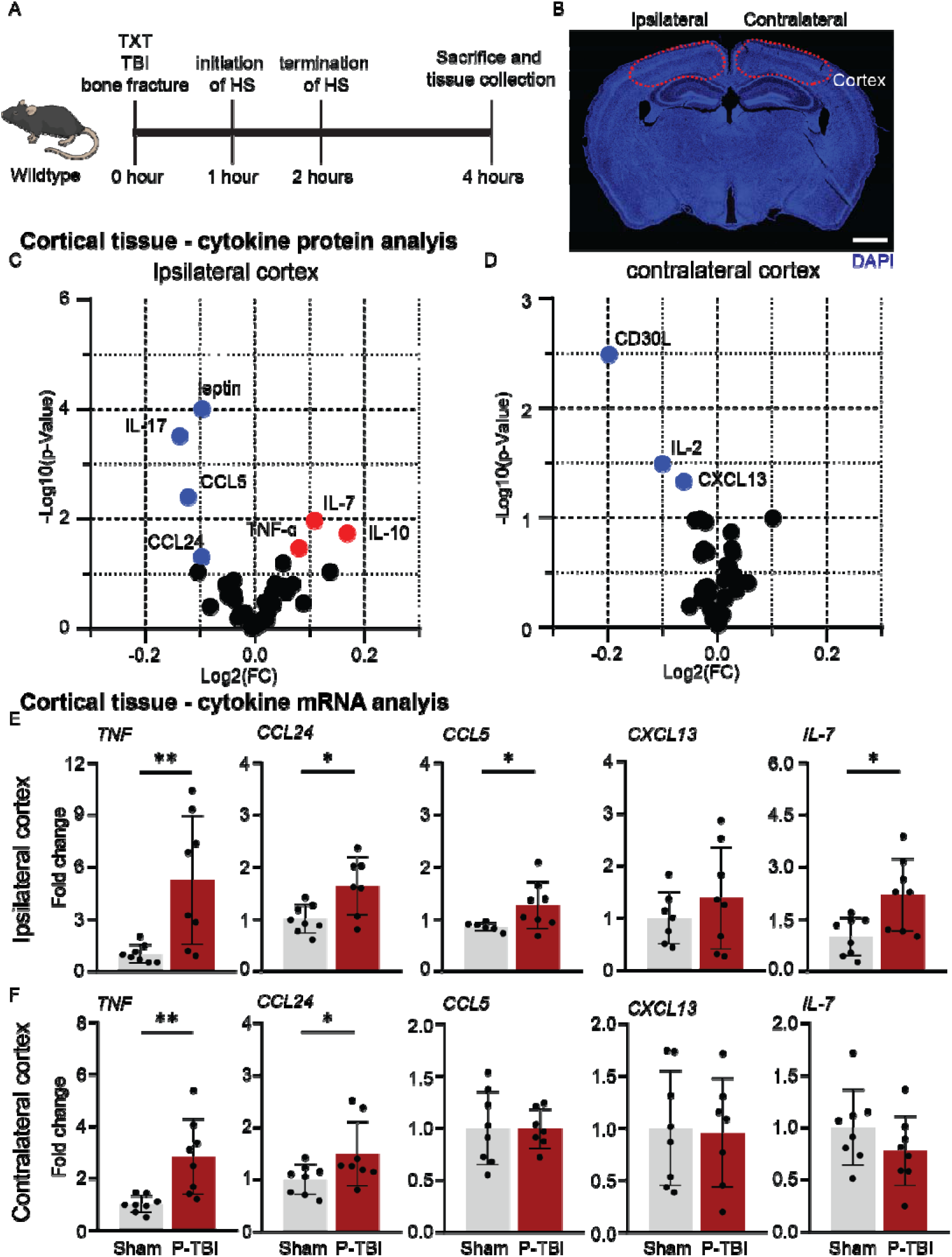
Polytrauma induces early cortical neuroinflammatory response. **A.** Experimental design of the polytrauma, consisting of blunt thorax trauma (TXT), traumatic brain injury (TBI), bone fracture and haemorrhagic shock (HS). The total observation time post P-TBI is 4 h followed by sacrifice and tissue collection. **B.** Representative image of DAPI stained in the mouse brain post P-TBI, highlighting the cortex. Scale bar: 200 um. **C-D.** Volcanoplot of cytokine protein analysis in cortical tissue revealed the significant upregulation of TNF-IZI, IL-7 and IL-10 and downregulation of CCL24, CCL5, IL-17 and leptin in the ipsilateral cortex. In the contralateral cortex significant downregulation of CXCL13, IL-2 and CD30L. **E-F.** Cytokine mRNA analysis in ipsilateral cortical tissue revealed the significant upregulation of TNF, CCL24, CCL5 and IL-7 (TNF: p=0.006, CCL24: p=0.014, CCL5: p=0.029, IL-7: p=0.013). The contralateral cortex revealed a significant increase of TNF and CCL24 (TNF-1.z: p=0.004, CCL24: p=0.049), with no difference in CCL5, CXCL13 and IL-7. Data shown as volcanoplots depicting the Log2 fold change and -Log10 p-value, and bar plots with individual data points. Group size: Sham N = 8, P-TBI N = 8. \**p* < 0.05; \*\**p* < 0.01.

As an entry point, we assessed the cytokine landscape in cortical samples subject to direct trauma (TBI-ipsilateral) and contralateral cerebral cortex from P-TBI (vs shams) using an antibody array covering 40 distinct inflammatory mediators (Fig.1C-D). Ipsilateral cortical samples displayed a significant elevation in the level of TNF, IL-7 and IL-10, and a downregulation of CCL5 and CCL24 (Eotaxin-2; Fig. 1C). IL-17 and Leptin were also significantly downregulated, but their absolute levels were very close to the detection limit. In contralateral cortical samples, we detected a downregulation of CXCL13 and CD30L (Fig. 1D; IL-2 was downregulated but at very low absolute levels).

We further verified these early signs of neuroinflammation associated with P-TBI by assessing the mRNA profiles of the immune mediators identified by the protein-based antibody array. We detected the substantial upregulation of *TNF* mRNA not only in ipsilateral samples (up to 12-fold the baseline; p = 0.0062; Fig. 1E) but also in the contralateral cortex (up to 5 times the baseline; p = 0.004; Fig. 1F). *IL-7* mRNA was also upregulated but only in the ipsilateral cortex (p = 0.013), in agreement with the corresponding protein levels (Fig. 1E-F). On the other hand, mRNA levels of *CCL24*/*Eotaxin-2*, *CCL5* were upregulated in the ipsilateral cortex (*CCL24/Eotaxin-2* mRNA also in the contralateral cortex, ipsi: p = 0.014, contra: p = 0.049; *CCL5* ipsi: p = 0.029; Fig. 1E-F), opposite to the protein dynamics. No change in mRNA levels of *CXCL13* were detected in ipsi- or contralateral cortical samples (Fig. 1E-F).

Thus, P-TBI results in early and diffuse (ipsilateral/contralateral) neuroimmune activation as highlighted by the upregulation of TNF mRNA and protein level.

### 2. Focal and diffuse neuroinflammation in acute TBI with polytrauma

The rapid and substantial elevation in *TNF* in both Ipsi and Contra cortical samples suggested the early induction of innate-immunity mediators. We confirmed this view by assessing the expression of a set of cytokines with established early/innate response roles, namely *CCL2, IL-1*β*, IL-33, IL-6* and the murine IL-8 homologues *CXCL1* and *CXCL2*. *CCL2* mRNA was revealed to be elevated both in ipsi- (Fig. 2A) and in contralateral (Fig. 2B) cortical samples, whereas *IL-1*β mRNA was upregulated only in the ipsilateral cortex (p = 0.002). *CXCL1* and *CXCL2* mRNA were significantly upregulated in the ipsilateral cortex (CXCL1: p < 0.0001; CXCL2: p = 0.006; Fig. 2A), but only *CXCL2* was also upregulated in the contralateral cortex (p = 0.011; Fig. 2B). No elevation was detected for *IL-33* and *IL-6*.

**Figure 2:**
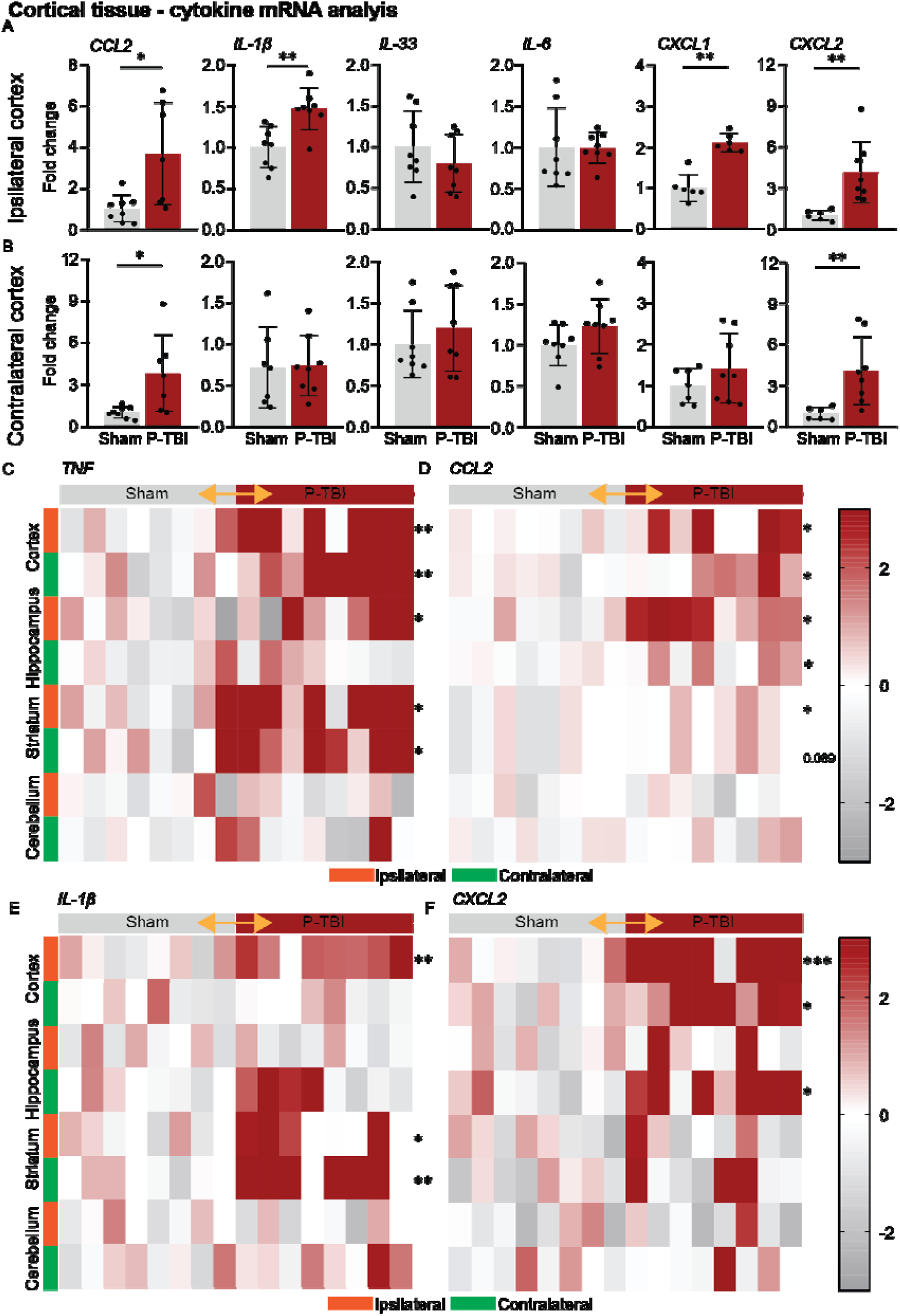
Focal and diffuse neuroinflammation in acute polytrauma with haemorrhagic shock. **A-B.** Cytokine mRNA analysis in ipsilateral cortical tissue revealed the significant increase of CCL2, IL-1□, CXCL1 and CXCL2 (CCL2: p=0.012, IL-1□: p=0.002, CXCL1: p<0.001, CXCL2: p=0.006). Contralateral cortical tissue revealed a significant increase of CCL2 and CXCL2 (CCL2: p=0.013, CXCL2: p=0.011). No significant difference in IL-33 and IL-6 was observed. **C-F.** Heatmaps depicting TNF, CCL2, IL-1β and CXCL2 expression in both sham vs P-TBI groups (comparison was made between sham and P-TBI; orange arrow, significance shown in stars next to the region) in ipsi- and contralateral cortex, hippocampus, striatum and cerebellum revealed large brain-wide increase of TNF (ipsi cortex: p=0.006, contra cortex: p=0.004, ipsi hippocampus: p=0.048, ipsi striatum: p=0.025, contra striatum: p=0.011) and CCL2 (ipsi cortex: p=0.012, contra cortex: p=0.013, ipsi hippocampus: p=0.013, contra hippocampus: p=0.02, ipsi striatum: p=0.022 contra striatum: p=0.069), in all regions except cerebellum. The upregulation of IL-1β was restricted to the ipsilateral cortex and both striata and the upregulation of CXCL2 in both cortici and contralateral hippocampus. Data shown as bar plots with individual data points and heatmaps depicting z-scores. Group size: Sham N = 8, P-TBI N = 8. \**p* < 0.05; \*\**p* < 0.01; ***p< 0.001.

We capitalized on the upregulation of *TNF*, *CCL2* and *IL-1*β upon P-TBI to explore the extent of spatial cerebral involvement in P-TBI, quantifying the expression of these cytokines in samples from cortex, hippocampus, striatum and cerebellum from ipsi- and contralateral sides. Substantial elevation of *TNF* mRNA was detected not only in ipsi- and contralateral cortex (p = 0.006; p = 0.004), but also in ipsilateral and contralateral striatum (p = 0.025; p = 0.011) and ipsilateral hippocampus (p = 0.048), though not in cerebellum, indicating a brain-wide induction (Fig. 2C). Likewise, *CCL2* upregulation was observed in ipsi- and contralateral cortex (p = 0.012; p = 0.013), hippocampus (p = 0.013; p = 0.02) and in striatum (p = 0.022; p = 0.069), but not in cerebellum (Fig. 2D). Conversely, *IL-1*β mRNA was increased only in the ipsilateral cortex (p = 0.002) as well as in striatum (bilaterally; p = 0.017; p = 0.002) but not in hippocampus or cerebellum (Fig. 2E). Notably, a significant *CXCL2* upregulation was detected only in cortex (bilaterally; p = 0.001; p = 0.017; Fig. 2F) and in the contralateral hippocampus (p = 0.014; Fig. 2F), but not in striatum or cerebellum.

Thus, in the early P-TBI response, a structure-specific and site-specific inflammatory milieu may develop, with *TNF* and *CCL2* upregulated throughout the brain, *IL-1*β overlapping with *TNF* only in cortex and striatum and *CXCL2* overlapping only in cortex.

Taken together, these findings highlight the co-occurrence of a focal, intense innate immunity activation, in correspondence of the ipsilateral cortex and a widespread, lower-intensity activation occurring across multiple ipsi- and contralateral brain structures.

### 3. Loss of TNF, CCL2 induction and reactive microglia in mice lacking C3 and CD14

Complement factor 3 (C3) and CD14 have been recognized as critical mediator of the early systemic innate immunity response involved in secondary TBI [17,18], in polytrauma [5] and in sepsis [19,20], with their roles often redundant. We therefore explored the effect of the combined loss of C3 and CD14 on the early brain response to P-TBI. To this aim, we subjected either wild-type mice or *C3^−/−^CD14^−/−^* double-knockout mice to P-TBI, obtained brain samples at the 4h timepoint and subjected them to single-molecule hybridization for *TNF* or *CCL2* mRNA (Fig. 3A-B). We included the immunolabeling of microglia (using the IBA1 antigen), since we hypothesized that these cells may strongly contribute to the early inflammatory response and be sensitive to C3 and/or CD14 loss.

**Figure 3:**
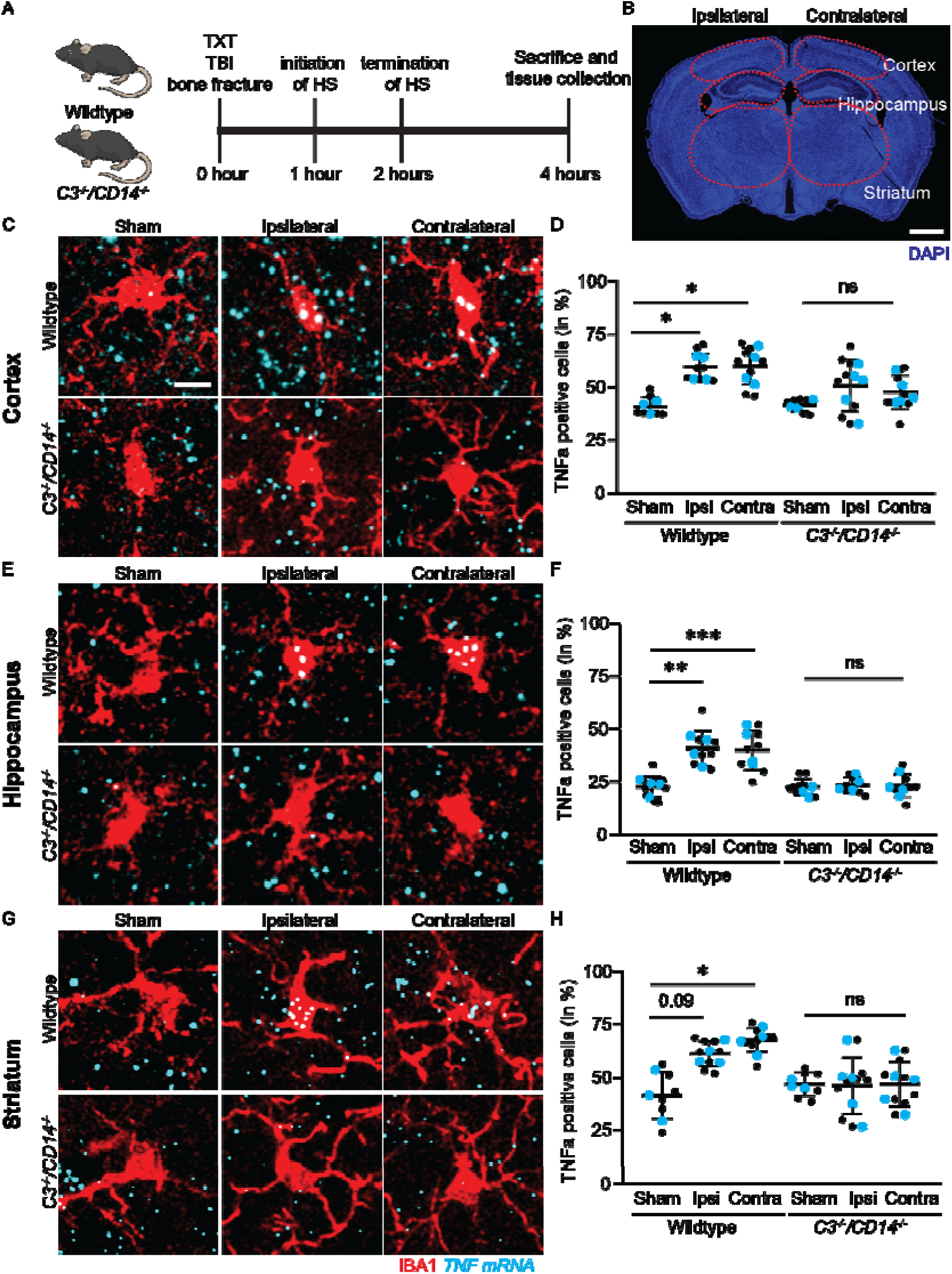
Brainwide Loss of P-TBI induced *TNF* expression in microglia of C3^−/−^/CD14^−/−^ mice. **A-B.** Experimental design of polytrauma and TBI (P-TBI) in wildtype and *C3^−/−^/CD14^−/−^* mice, investigating ipsilateral and contralateral cortex, hippocampus and striatum. **C-H.** Single mRNA *in situ* hybridization of TNF in IBA1+ microglia revealed significant increase of TNF-1.z+ microglia in ipsi- and contralateral cortex (WT sham vs WT P-TBI ipsi: p = 0.021; WT sham vs WT P-TBI contra: p = 0.015), hippocampus (WT sham vs WT P-TBI ipsi: p = 0.002; WT sham vs WT P-TBI contra: p = 0.001) and striatum (WT sham vs WT P-TBI ipsi: p = 0.09; WT sham vs WT P-TBI contra: p = 0.017) of wildtype mice. No increase of TNF+ microglia was observed in bilateral cortex, hippocampus or striatum for *C3^−/−^/CD14^−/−^*mice. Data shown as dot plots with all data points depicted and single mice as coloured dots (cyan). Scale bar: 10 um. Group size: WT sham N = 4, WT P-TBI N = 4. C3^−/−^/CD14^−/−^ Sham N = 3, C3^−/−^/CD14^−/−^ mice P-TBI N = 4. \**p* < 0.05; \*\**p* < 0.01; ***p< 0.001; ns = not significant.

First, we quantified the overall density of *TNF* mRNA (number of molecules per surface unit, irrespective of their cellular source). Confirming the qPCR, we noted an increase in the expression of *TNF* mRNA in the cortex (ipsi: p = 0.007; contra: p = 0.017; Suppl. Fig. 1D, E), hippocampus (ipsi: p = 0.001; contra: p = 0.01; Suppl. Fig. 1, F) and striatum (ipsi: p = 0.007; contra: p = 0.01; Suppl. Fig. 1D, G) of WT mice, in both ipsilateral and contralateral samples. Coherently, we also detected a significant increase in microglial cells expressing *TNF* mRNA, i.e. containing at least 4 mRNA molecules detected by the in-situ probe, in cortex (ipsi: p = 0.021; contra: p = 0.015; Fig. 3C, D), hippocampus (ipsi: p = 0.002; contra: p = 0.001; Fig. 3E, F) and a strong trend in striatum (ipsi: p = 0.093; contra: p = 0.017; Fig. 3G, H) of WT mice subjected to P-TBI compared to sham controls, again with minimal differences between ipsi- and contralateral side. These findings confirmed the upregulation of *TNF* mRNA and identified microglia as a prominent source.

Most notably, the increase in *TNF* mRNA was abrogated in the contralateral side of the *C3^−/−^CD14^−/−^* in cortex (ipsi: p = 0.045; contra: p = 0.092; Suppl. Fig. 1D, E), hippocampus (ipsi: p = 0.048; contra: p = 0.893; Suppl. Fig. 1D, F) and striatum (ipsi: p = 0.037; contra: p = 0.741; Suppl. Fig. 1D, G). When looking at the expression in IBA1+ microglia upon P-TBI, *TNF* increase was completely abrogated in the *C3^−/−^CD14^−/−^* mice, in cortex (ipsi: p = 0.537; contra: p = 0.697; Fig. C, D), hippocampus (ipsi: p = 0.865; contra: p = 0.959; Fig. 3E, F) and in the striatum (ipsi: p > 0.999; contra: p > 0.999; Fig. 3G, H).

Second, we confirmed the *TNF* data by assessing the quantity and location of the *CCL2* mRNA. We found a substantial expression of *CCL2* in microglia, and the number of IBA1+/*CCL2* mRNA+ cells significantly increased in WT animals subject to P-TBI, again across the anatomical spectrum of either ipsi- or contralateral cortex (ipsi: p = 0.009; contra: p = 0.012; Fig. 4A, B), hippocampus (ipsi: p = 0.004; contra: p = 0.009; Fig. 4C, D) or striatum (ipsi: p = 0.003; contra: p = 0.011; Fig. 4E, F). On the other hand, no such increase in IBA1+ cells expressing *CCL2* mRNA+ was observed in *C3^−/−^CD14^−/−^*across cortex (ipsi: p = 0.479; contra: p = 0.668; Fig. 4A, B), hippocampus (ipsi: p = 0.458; contra: p = 0.739; Fig. 4C, D) nor striatum (ipsi: p = 0.888; contra: p = 0.993; Fig. 4E, F).

**Figure 4:**
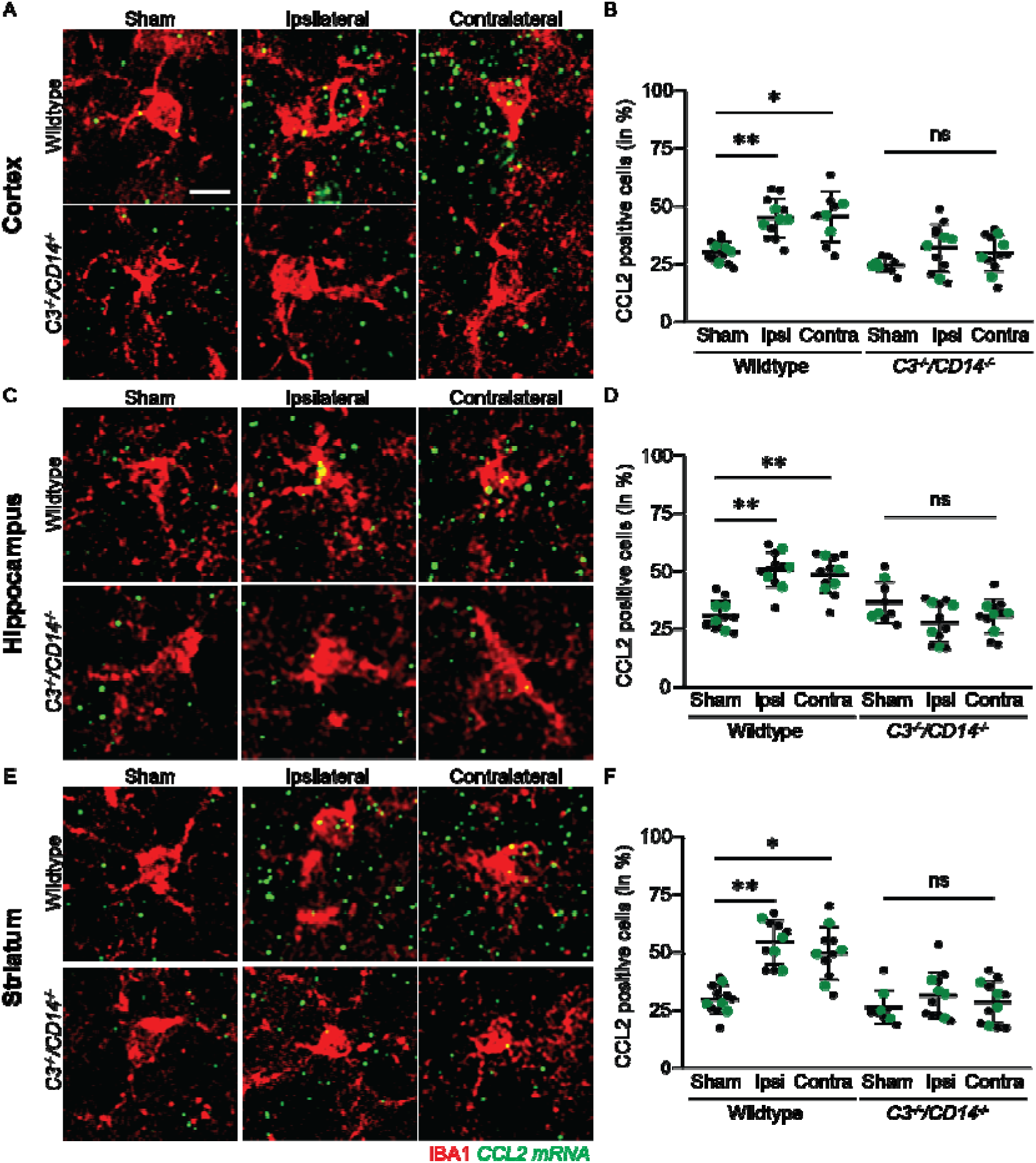
Brain wide Loss of P-TBI induced *CCL2* expression in microglia of C3^−/−^/CD14^−/−^ mice. **A-F.** Single mRNA *in situ* hybridization of CCL2 in IBA1+ microglia revealed significant increase of CCL2+ microglia in ipsi- and contralateral cortex (WT sham vs WT P-TBI ipsi: p = 0.009; WT sham vs WT P-TBI contra: p = 0.012), hippocampus (WT sham vs WT P-TBI ipsi: p = 0.004; WT sham vs WT P-TBI contra: p = 0.009) and striatum (WT sham vs WT P-TBI ipsi: p = 0.003; WT sham vs WT P-TBI contra: p = 0.011) of wildtype mice. No increase of TNF+ microglia was observed in bilateral cortex, hippocampus or striatum for C3^−/−^/CD14^−/−^ mice. Data shown as dot plots with single sections depicted as black dots and single mice as coloured dots (green). Scale bar: 10 um. Group size: WT sham N = 4, WT P-TBI N = 4. C3^−/−^/CD14^−/−^ Sham N = 3, C3^−/−^/CD14^−/−^ mice P-TBI N = 4. \**p* < 0.05; \*\**p* < 0.01; ***p< 0.001; ns = not significant.

Third, we sought to confirm the effect of *C3^−/−^CD14^−/−^*on the reactivity of microglia to P-TBI across the brain, using the phosphorylation of the ribosomal protein S6 as marker of cellular response. When phosphorylated, S6 enhances protein synthesis and S6 phosphorylation is used to identify cells undergoing active protein synthesis in response to stimuli [21,22]. Immunolabelling intensity for pS6 in IBA1+ cells substantially increased in WT animals subject to P-TBI, compared to sham, in ipsi-as well as in contralateral cortex (ipsi: p = 0.008; contra: p = 0.007; Fig. 5C, D), again with limited difference between the two sides. Likewise, a significant increase in pS6 immunoreactivity in microglia was observed in ipsi- and contralateral hippocampus (ipsi: p = 0.036; contra: p = 0.013; Fig. 5E, F) and in ipsilateral striatum (ipsi: p = 0.004; contra: p = 0.088; Fig. 5G, H). Notably, also S6 phosphorylation induced by P-TBI in microglia was completely abolished in *C3^−/−^CD14^−/−^*mice.

**Figure 5:**
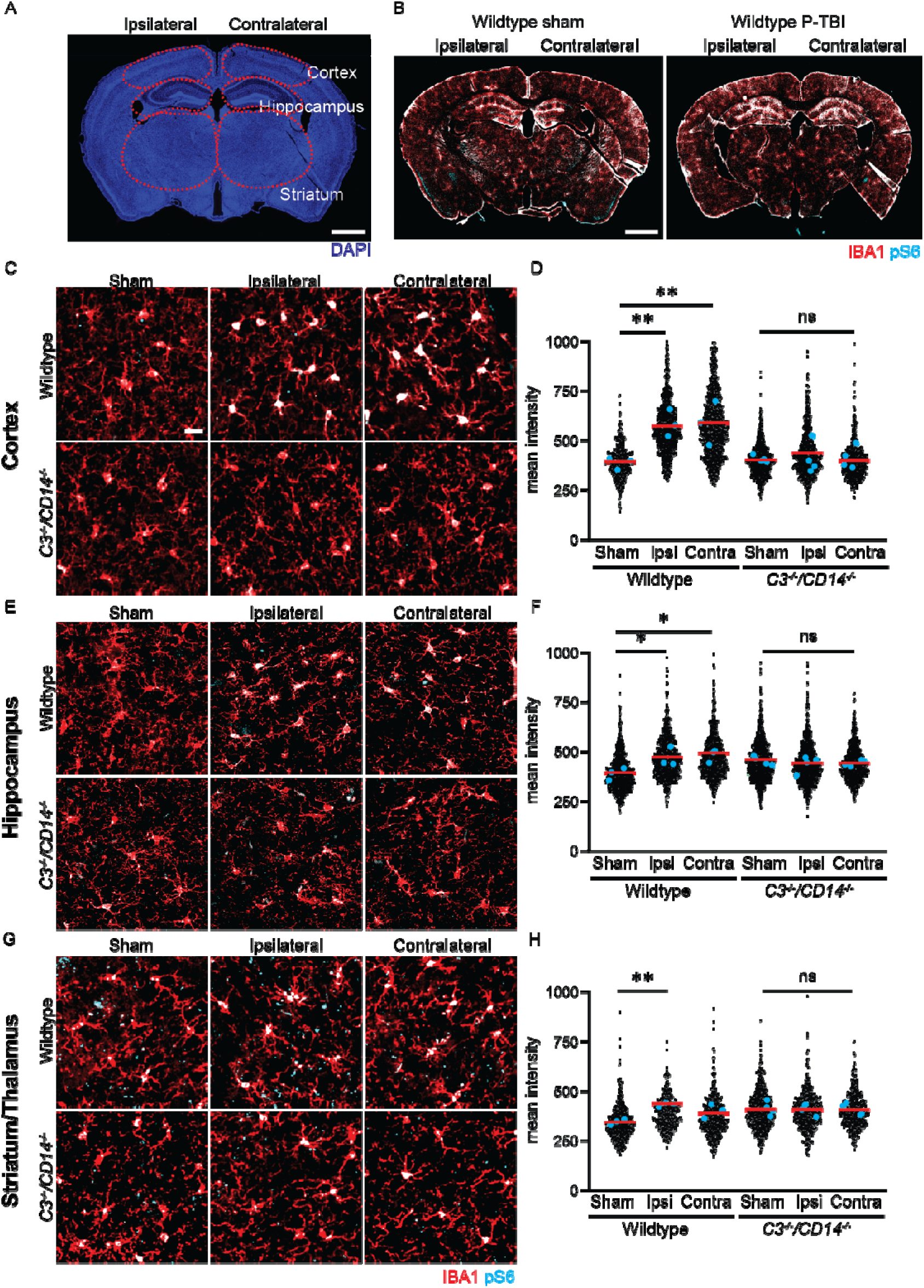
Loss of P-TBI induced microglia activity in C3^−/−^/CD14^−/−^ mice. **A-B.** Representation of analyzed brain regions of wildtype mice post sham and P-TBI, depicting a brainwide induction of pS6 in IBA1+ microglia post P-TBI. **C-H.** Immunofluorescence staining and imaging of pS6 and IBA1 revealed increase of pS6 intensity in microglia in both ipsi- and contralateral cortex (WT sham vs WT P-TBI ipsi: p = 0.008; WT sham vs WT P-TBI contra: p = 0.007), hippocampus (WT sham vs WT P-TBI ipsi: p = 0.036; WT sham vs WT P-TBI contra: p = 0.013) and striatum (WT sham vs WT P-TBI ipsi: p = 0.004; WT sham vs WT P-TBI contra: p = 0.088). No increase of pS6 intensity was observed in bilateral cortex, hippocampus or striatum for *C3^−/−^/CD14^−/−^*mice. Data shown as dot plots with single microglia depicted as black dots and single mice as coloured dots (cyan). Scale bar overview: 200 um; insert: 10 um. Group size: WT sham N = 4, WT P-TBI N = 4. C3^−/−^/CD14^−/−^ Sham N = 3, C3^−/−^/CD14^−/−^ mice P-TBI N = 4. \**p* < 0.05; \*\**p* < 0.01; ***p< 0.001; ns = not significant.

In addition to the expression and phosphorylation readouts, we also examined the morphology of IBA1+ microglia, since reactive microglia is recognized to lose its ramified morphology in favour of an ameboid appearance (Fig. 6A-B; [23]). Whereas microglia from WT and *C3^−/−^CD14^−/−^*displayed a similar ramified architecture in sham animals (as quantified in the Sholl analysis; Fig. 6C-D), upon P-TBI, microglia in WT animals displayed a significant decrease in ramification compared to *C3^−/−^CD14^−/−^*(thus, shifting toward an ameboid morphology) in the bilateral cortex (ipsi WT P-TBI vs *C3^−/−^CD14^−/−^ P-TBI*: p < 0.05 at 8-16, 19, 25-27 and 32-36 um; contra WT P-TBI vs *C3^−/−^CD14^−/−^ P-TBI*: p < 0.05 at 6 and 29 um; Fig. 6E-G) and bilateral hippocampus (ipsi WT P-TBI vs *C3^−/−^CD14^−/−^ P-TBI*: p < 0.05 at 19-30 um; contra WT P-TBI vs *C3^−/−^CD14^−/−^ P-TBI*: p < 0.05 at 19-25 and 29 um; Fig. 6H-J), though no change was observed in the striatum (Fig. 6K-M).

**Figure 6:**
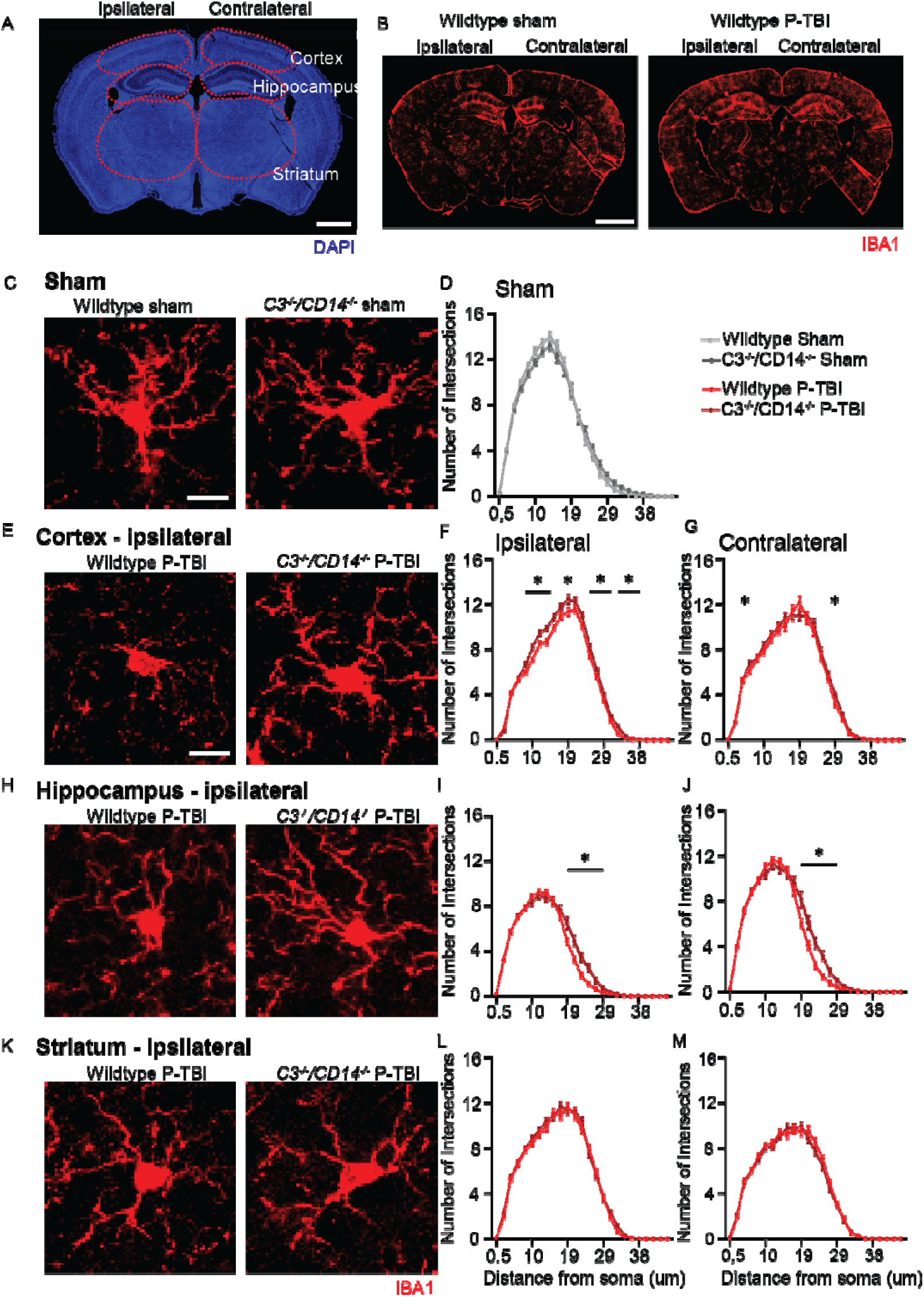
Loss of P-TBI induced reactive microglia morphology in C3^−/−^/CD14^−/−^ mice. **A-B.** Representation of analyzed brain regions of wildtype mice post sham and P-TBI, showing a brainwide induction IBA1+ microglia post P-TBI. **C-D.** Immunofluorescence staining and imaging of IBA1+ microglia morphology revealed no differences between WT and *C3^−/−^/CD14^−/−^* sham. **E-M.** Immunofluorescence staining and imaging of IBA1+ microglia morphology revealed a decrease in the number of intersections between wildtype and *C3^−/−^/CD14^−/−^* mice post P-TBI in ipsilateral cortex (WT P-TBI ipsi vs *C3^−/−^/CD14^−/−^* P-TBI ipsi: p < 0.05 at 8-16, 19, 25-27 and 32-36 um) as well as ipsi- and contralateral hippocampus (WT P-TBI ipsi vs *C3^−/−^/CD14^−/−^*P-TBI ipsi: p < 0.05 at 19-30 um; WT P-TBI contra vs *C3^−/−^/CD14^−/−^*P-TBI contra: p < 0.05 at 19-25 and 29 um). No difference was observed in striatum. Data shown as X-Y plots depicting number of intersections over distance from soma. Scale bar overview: 200 um; 10 um. Group size: n = 80 - 120 microglia per treatment group; WT P-TBI N = 4, C3^−/−^/CD14^−/−^ P-TBI N = 4. \**p* < 0.05.

Taken together, these findings show that microglia across cortex, hippocampus and striatum react to P-TBI with an increased transcription of TNF and CCL2 mRNA, an upregulation of protein synthesis and a shift toward an ameboid morphology. Of note, all these manifestations of the reactive phenotype are abolished in animals lacking C3 and CD14.

### 4. Induction of TNF and CCL2 upon P-TBI depends on C3 in cortex and striatum

Although C3 and CD14 usually reveal a converging role in driving inflammatory responses [10,24], several instances are known in which their effects diverge or turn anti-inflammatory (e.g., CD14 is anti-inflammatory in the context of efferocytosis [25] whereas the C3 fragment iC3b suppresses proinflammatory cytokines from macrophages [26]). Since the double *C3^−/−^CD14^−/−^*knock-out did not allow the disentanglement of any differential effect of C3 vs CD14 in the early neuroimmune activation caused by P-TBI, we considered an in-depth investigation involving four distinct mouse genotypes : WT, single-KO *C3^−/−^*, single-KO *CD14^−/−^* and double-KO *C3^−/−^CD14^−/−^*, each group divided in subgroups subjected to P-TBI or sham (altogether, 8 treatment x genotype groups; Fig. 7A). At the beginning, we verified that C3 mRNA was undetectable in *C3^−/−^*and *C3^−/−^CD14^−/−^* animals whereas CD14 mRNA was undetectable in *CD14^−/−^* and *C3^−/−^CD14^−/−^*animals (Fig. 7C-D).

**Figure 7:**
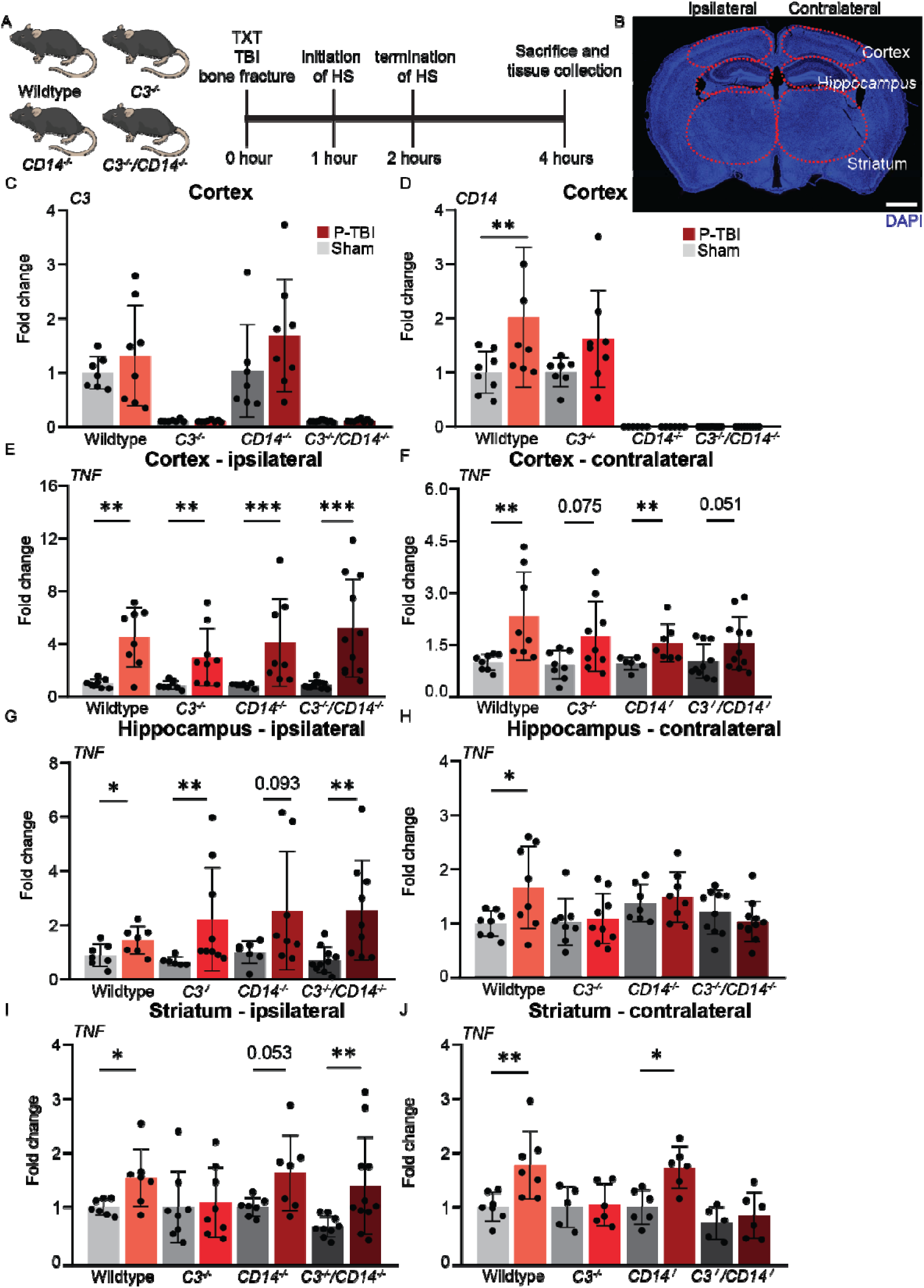
P-TBI induced *TNF* expression in the cortex and striatum depends on C3. **A-B.** Experimental design of polytrauma and TBI (P-TBI) in wildtype, *C3^−/−^*, *CD14^−/−^*and *C3^−/−^/CD14^−/−^* mice, investigating bilateral cortex, hippocampus and striatum. **C-D.** Real-time quantitative PCR (RT-PCR) of *C3* and *CD14* expression in cortical tissue validates wildtype, *C3^−/−^*, *CD14^−/−^* and *C3^−/−^/CD14^−/−^*mice. **E-F.** RT-PCR analysis of *TNF* expression in ipsi- and contralateral cortex post sham or P-TBI revealed a significant increase in all genotypes in the ipsilateral cortex (Sham vs P-TBI; WT: p = 0.005; *C3^−/−^*: p = 0.006; *CD14^−/−^*: p = 0.0003; *C3^−/−^/CD14^−/−^*: p < 0.0001). In the contralateral cortex an overall increase was observed, with significance only in WT and *CD14^−/−^* mice (Sham vs P-TBI; WT: p = 0.001; *C3^−/−^*: p = 0.075; *CD14^−/−^*: p = 0.009; *C3^−/−^/CD14^−/−^*: p = 0.051). **G-H.** RT-PCR analysis of *TNF* expression in ipsi- and contralateral hippocampus post sham or P-TBI revealed a significant increase in all genotypes, except *CD14^−/−^* in the ipsilateral hippocampus (Sham vs P-TBI; WT: p = 0.046; *C3^−/−^*: p = 0.002; *CD14^−/−^*: p = 0.094; *C3^−/−^/CD14^−/−^*: p = 0.004). In the contralateral hippocampus a significant increase was observed only in the WT (Sham vs P-TBI; WT: p = 0.033). **I-J.** RT-PCR analysis of *TNF* expression in ipsi- and contralateral striatum post sham and P-TBI revealed a significant increase in WT and *C3^−/−^/CD14^−/−^*in the ipsilateral striatum (Sham vs P-TBI; WT: p = 0.025; *CD14^−/−^*: p = 0.053; *C3^−/−^/CD14^−/−^*: p = 0.01). In the contralateral striatum a significant increase was observed only in the wildtype and *CD14^−/−^* (Sham vs P-TBI; wildtype: p = 0.007; CD14^−/−^: p = 0.015). Data shown as bar plots with individual data points. Group size: WT sham N = 8, WT P-TBI N = 8, C3^−/−^ sham N = 8, C3^−/−^P-TBI N = 8, CD14^−/−^ sham N = 7, CD14^−/−^ P-TBI N = 8, C3^−/−^/CD14^−/−^ sham N = 10, C3^−/−^/CD14^−/−^ P-TBI N = 11. \**p* < 0.05; \*\**p* < 0.01; ***p< 0.001.

First, we considered the induction of *TNF* mRNA (Fig.3C-D) upon P-TBI across the four genotypes vs sham controls for each genotype, examining ipsi and contralateral samples from cortex, hippocampus and striatum. Interestingly, in ipsilateral cortical samples (the area subject to the highest mechanical stress; Fig. 7B), P-TBI upregulated *TNF* mRNA but neither the loss of C3, nor of CD14 nor their combined loss significantly changed *TNF* induction (in agreement with the single-molecule data; Fig. 7E). Likewise, in contralateral cortical samples P-TBI upregulated *TNF* mRNA (p = 0.001), with a non-significant trend remaining upon loss of C3 and combined C3 and CD14 (p = 0.075; p = 0.051, respectively), but a strong significant increase still persistent upon loss of CD14 (p = 0.009; Fig. 7F). WT hippocampal samples also exhibited a robust induction of *TNF* in ipsilateral, which was largely unaffected in *C3^−/−^* and/or *CD14^−/−:^*animals (WT: p = 0.046; *C3^−/−^*: p = 0.002; *CD14^−/−:^*: p = 0.094; *C3^−/−^CD14^−/−^*: p = 0.004; Fig. 7G), whereas in contralateral samples with either C3*^−/−^*, CD14*^−/−^*or combined loss suppressed the *TNF* response (WT: p = 0.033; Fig. 7H). In the striatum, P-TBI caused a comparable upregulation of *TNF* in both ipsi- and contralateral samples of WT mice (ipsi: p = 0.025; contra: p = 0.007). On the ipsilateral side, this effect was completely abolished in *C3^−/−^*animals (but not in *CD14^−/−^* or *C3^−/−^CD14^−/−^*; p = 0.053 and 0.01 respectively; Fig. 7I) whereas in contralateral samples, *TNF* induction was completely abolished in both *C3^−/−^* and in *C3^−/−^CD14^−/−^*animals (though not *CD14^−/−^*: p = 0.015; Fig. 7J).

Thus, *TNF* mRNA induction in the contralateral cortex and striatum is dependent on C3 whereas hippocampal *TNF* induction shows sensitivity to both C3 and CD14. In the ipsilateral cortex and hippocampus, the induction of *TNF* mRNA is not dependent on either C3 or CD14, whereas it appears dependent on C3 in the ipsilateral striatum.

We sought an independent confirmation of the region-specific roles of C3 and CD14 in the neuroinflammatory response to P-TBI by assessing the expression of *CCL2* across genotypes and brain structures. Surprisingly, *CCL2* expression in the ipsilateral cortex was moderately increased in WT samples but was actually enhanced in *C3^−/−^* and *C3^−/−^CD14^−/−^*animals (WT: p = 0.012; *C3^−/−^*: p = 0.01; *CD14^−/−^*: p = 0.023 or *C3^−/−^CD14^−/−^*: p < 0.0001; Fig. 8A). *CCL2* induction in WT contralateral cortex (p = 0.031) was, on the contrary, suppressed upon the loss of C3 and CD14 (*C3^−/−^, CD14^−/−^*and *C3^−/−^CD14^−/−^* animals; Fig. 8B). Upregulation of *CCL2* in the ipsilateral hippocampus was detected in WT samples as well as in animals lacking C3 and/or CD14 (WT: p = 0.013; *C3^−/−^*: p = 0.001; *CD14^−/−^*: p = 0.04 or *C3^−/−^CD14^−/−^*: p = 0.001; Fig. 8C), whereas *CCL2* upregulation in the contralateral hippocampus was suppressed by the loss of either C3, or CD14 or both (Fig. 8D). The induction of *CCL2* in the ipsilateral striatum was modest in magnitude (p = 0.022) but appeared to be completely lost in *C3^−/−^* but not in *CD14^−/−^* or *C3^−/−^CD14^−/−^* animals (p = 0.019 and 0.043 respectively); conversely, induction of *CCL2* in the contralateral striatum was not significant (Fig. 8E-F). This data is in accordance with the *in situ* hybridization data of *CCL2* expression of WT and *C3^−/−^CD14^−/−^* in cortex, hippocampus and striatum (Suppl. Fig. 1 H-K).

**Figure 8:**
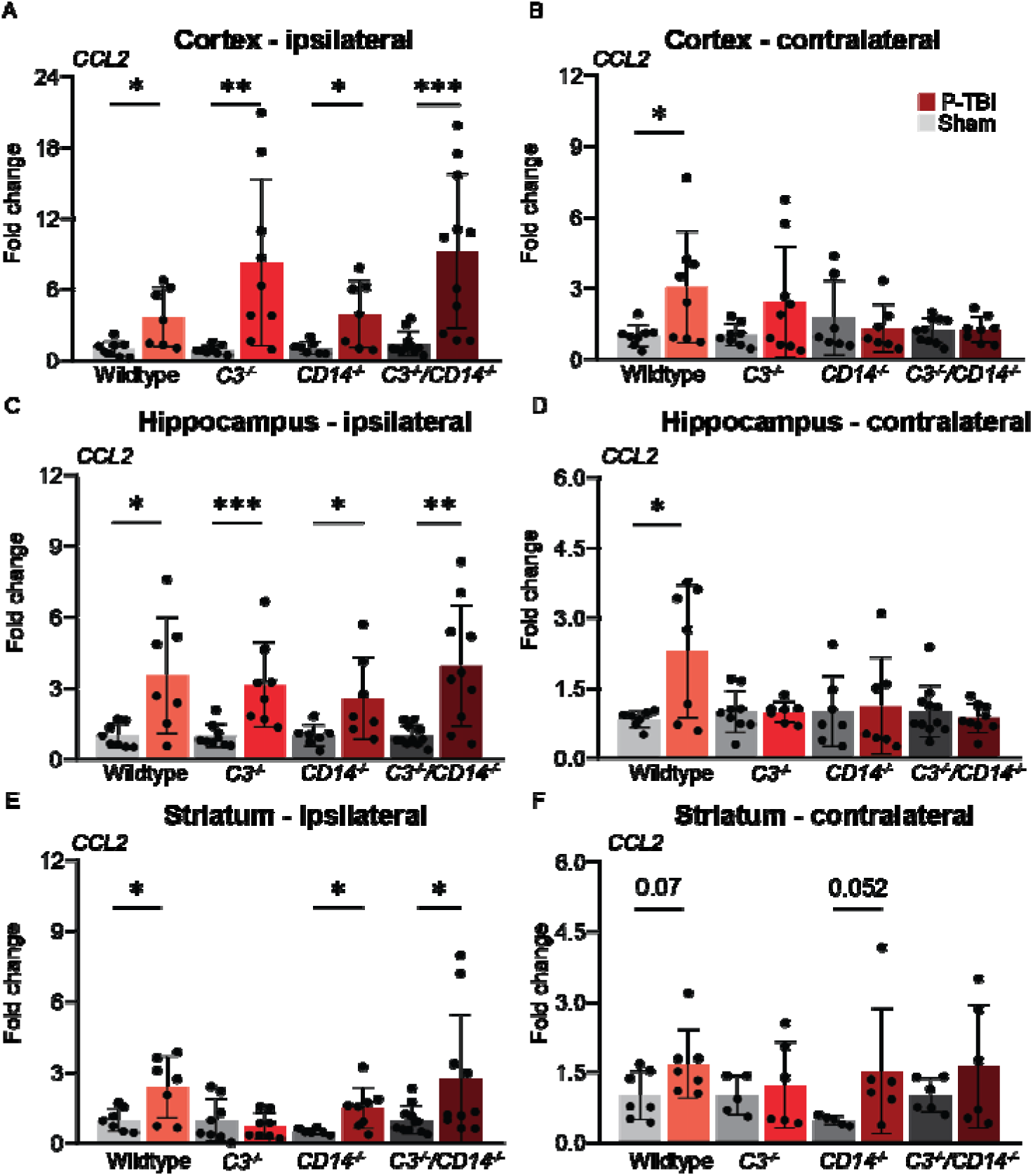
P-TBI induced *CCL2* expression in the cortex depends on CD14, and in the striatum depends on C3. **A-B.** RT-PCR analysis of *CCL2* expression in ipsi- and contralateral cortex post sham and P-TBI revealed a significant increase in all genotypes in the ipsilateral cortex (Sham vs P-TBI; WT: p = 0.012; *C3^−/−^*: p = 0.01; *CD14^−/−^*: p = 0.023; *C3^−/−^/CD14^−/−^*: p < 0.0001). In the contralateral cortex a significant increase was observed only in WT, with strong trend in C*3^−/−^* (Sham vs P-TBI; WT: p = 0.031; *C3^−/−^*; p = 0.37). **C-D.** RT-PCR analysis of *CCL2* expression in ipsi- and contralateral hippocampus post sham and P-TBI revealed a significant increase in all genotypes in the ipsilateral hippocampus (Sham vs P-TBI; WT: p = 0.013; *C3^−/−^*: p = 0.001; *CD14^−/−^*: p = 0.04; *C3^−/−^/CD14^−/−^*: p = 0.001). In the contralateral hippocampus a significant increase was observed only in WT (Sham vs P-TBI; WT: p = 0.02). **E-F.** RT-PCR analysis of *CCL2* expression in ipsi- and contralateral striatum post sham and P-TBI revealed a significant increase in WT, *CD14^−/−^* and *C3^−/−^/CD14^−/−^*in the ipsilateral striatum (Sham vs P-TBI; WT: p = 0.022; *CD14^−/−^*: p = 0.019; *C3^−/−^/CD14^−/−^*: p = 0.004). In the contralateral striatum a strong trend was observed only in the WT and *CD14^−/−^* (Sham vs P-TBI; WT: p = 0.069; *CD14^−/^*^−^: p = 0.052). Data shown as bar plots with individual data points. Group size: WT sham N = 8, WT P-TBI N = 8, C3^−/−^ sham N = 8, C3^−/−^ P-TBI N = 8, CD14^−/−^ sham N = 7, CD14^−/−^ P-TBI N = 8, C3^−/−^/CD14^−/−^ sham N = 10, C3^−/−^/CD14^−/−^ P-TBI N = 11. \**p* < 0.05; \*\**p* < 0.01; ***p< 0.001.

Thus, the *CCL2* response to P-TBI displays similarities to *TNF* with regards to C3 dependency in the contralateral hippocampus and striatum, with difference in the contralateral cortex, where the *CCL2* induction is largely CD14 dependent. For the ipsilateral site, a dependency of C3 was observed in the striatum only where the loss of CD14 resulted in a pro-inflammatory response that overcame the C3 loss.

### 5. C3 and CD14 do not affect systemic haemodynamic or organ damage in P-TBI

Having observed the differential effect of C3 and CD14 on the induction of a cerebral neuroinflammatory response in P-TBI, we reasoned that the central effects may actually be explained by differences in systemic inflammatory responses and peripheral organ damage. Thus, we considered the haemodynamic profiles in WT, *C3^−/−^*, *CD14^−/−^* and in *C3^−/−^CD14^−/−^*animals during the HS/resuscitation phase of the P-TBI (Suppl. Figure 2A). Plasma levels of C3b confirmed the genotype of mice, with no differences upon P-TBI (Suppl. Fig. 2B). Arterial pressure trends were comparable across the four genotypes in both sham groups as well as P-TBI groups (Suppl. Fig. 2C-E). As expected, heart rate showed a significant increase at around 180 minutes into the procedure between sham and P-TBI (WT sham vs WT P-TBI: p < 0.05 at 155, 180-185 and 195-205 min; Suppl. Fig. 2F). When considering the different genotypes, a small significant increase was observed between WT and *CD14^−/−^* mice around 180 min into the procedure in both sham and P-TBI groups (WT sham vs *CD14^−/−^*sham: p < 0.05 at 180, 195-205 min; WT P-TBI vs *CD14^−/−^* P-TBI: p < 0.05 at 180-185 and 195-205 min; Suppl. Fig. 2G-H).

Furthermore, we explored a battery of plasma damage markers related to specific organs, including brain (astroglial, GFAP), lungs (CC16), liver (ALT), pancreas (Lipase), intestines (I-FABP) and vasculature (syndecan), or systemic inflammatory markers, like: PAI-1, IL-6 and C5a. All damage markers were significantly increased upon P-TBI in WT mice (Suppl. Fig. 2I-Q), but were largely unaffected by the loss of C3 and/or CD14: no effect was found in glial damage, liver damage, kidney damage and systemic inflammation (C5a, IL-6) and only isolated effects on the intestine (I-FABP in *CD14^−/−^* and *C3^−/−^CD14^−/−^* ; Suppl. Fig. 2M) and the vascular system (syndecan in *C3^−/−^*; Suppl. Fig. 2N).

Thus, the absence of C3 and/or CD14 does not result in significant improvement in hemodynamic parameters, tissue damage or inflammation upon P-TBI, strongly suggesting that the observed differences at the cerebral level are due to focal alterations in the neuroinflammatory response rather than consequences of changes in systemic inflammation.

### 6. The expression pattern of C3 and CD14 across brain structures upon P-TBI accounts for the differential requirement for the induction of inflammatory cytokines

In order to explain the differential effect of C3 and CD14 in the induction of *TNF* and *CCL2* in the cortex vs striatum vs hippocampus, we first investigated the expression of *C3* and *CD14* mRNA by qPCR. Expression of *C3* was very limited but revealed a striking pattern, being weakest in the cortex and strongest in the striatum (Fig. 9A). Conversely, *CD14* expression was substantially higher than *C3* in absolute terms, and it was comparable in the striatum and in the cortex but was substantially weaker in the hippocampus (Fig. 9B).

**Figure 9:**
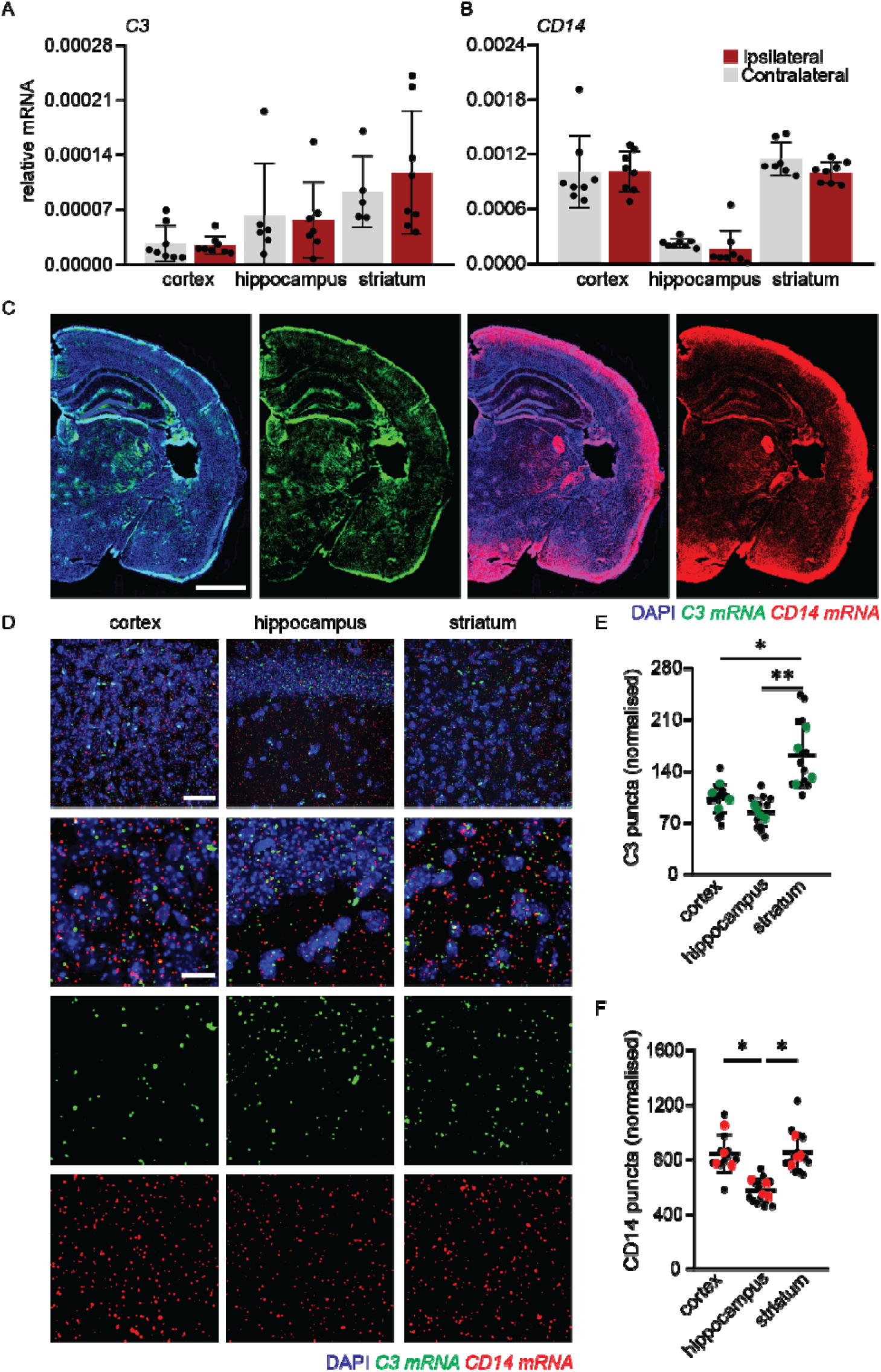
Differential pattern of C3 and CD14 expression across brain regions. **A-B.** RT-PCR analysis of *C3* and *CD14* expression in WT cortex, hippocampus and striatum revealed low, intermediate and high expression of *C3* in cortex, hippocampus and striatum respectively. For *CD14*, a high, low, high expression in cortex, hippocampus and striatum was observed. **C-F.** Single mRNA *in situ* hybridization of *C3* and *CD14* expression in cortex. *C3* showed highest expression in striatum, compared to cortex and hippocampus (cortex vs striatum: p = 0.005; hippocampus vs striatum: p = 0.011). *CD14* showed highest expression in cortex and striatum, and lowest expression in hippocampus (cortex vs hippocampus: p = 0.011; hippocampus vs striatum: p = 0.015). Data shown as bar plots with individual data points or dot plots as dot plots with single sections depicted as black dots and single mice as coloured dots (green/red). Scale bar overviews: 200 and 50 um; insert: 10 um. Group size: WT sham ipsi and contra expression data N = 8; RNAscope data N = 4. \**p* < 0.05; \*\**p* < 0.01.

We confirmed this expression pattern using single-molecule *in situ* hybridization. The density of *C3* mRNA was higher in the striatum than in the cortex or the hippocampus on average (cortex vs striatum: p = 0.005; hippocampus vs striatum: p = 0.011; Fig. 9D-E). However, *C3* mRNA distribution was not homogeneous: in the cortex, the expression was concentrated in the layer IV and V, where in the hippocampus it overlapped the neuronal outlines of dentate gyrus and CA fields (Fig. 9C). On the other hand, the density of *CD14* was comparable in the cortex and striatum (in agreement with qPCR data) and weaker in the hippocampus (cortex vs hippocampus: p = 0.011; hippocampus vs striatum: p = 0.015; Fig. 9D, F); *CD14* was diffusely expressed through the cortex and the striatum, whereas in the hippocampus was concentrated in the stratum oriens and stratum moleculare (Fig. 9C).

Thus, the expression data indicate that the pattern of dependency on C3 or CD14 for neuroinflammatory activation after P-TBI cannot be explained by their expression level. Both high-expressing striatum and low-expressing cortex rely on C3 for the development of diffuse neuroinflammation following P-TBI, even in the presence of robust CD14 expression.

## Discussion

Our investigation of the P-TBI model, including bone fracture, thorax trauma, HS and mild TBI, has revealed an early and diffuse induction of inflammatory cytokines across the brain, highlighted by the upregulation of *TNF*, *CCL2* and *IL-1*β and the appearance of reactive microglia. Contrary to our hypothesis, systemic inflammation and organ damage is not suppressed in mice lacking C3 or CD14, although in animals the cerebral inflammatory response is prevented or reduced and the microglial reactivity is abolished. Interestingly, the upregulation of inflammatory cytokines was differentially regulated by C3 more than by CD14. In structures with either high or low C3 expression (striatum and cortex), C3 was necessary to drive the expression of cytokines.

Our finding of a rather diffuse induction of inflammatory cytokines across the brain, on which the focal induction by TBI is superimposed, are cross-validated by previous findings showing a substantial degree of neuroinflammation occurring upon peripheral injury [27], polytrauma [28,29], or more broadly, during systemic inflammatory conditions [30] such as in sepsis [15]. In our model, as well as in previous reports [28], the cerebellum remains somehow unaffected. Interestingly, we identified a significant upregulation of *CCL2* in the cortex and hippocampus, but an actual downregulation was previously reported [28]; however, we considered an earlier timepoint (4h vs 6h) and our polytrauma was less severe (not including peripheral limb ischemia or amputation) than the previous one, suggesting that the neuroinflammatory response to P-TBI is context-dependent. Furthermore, the neuroinflammatory response after P-TBI is also structure-dependent: the response occurring in the striatum lacks the *CXCL2* upregulation observed in the cortex and hippocampus; this finding is in agreement with the distinct segregation of striatal samples in previously reported P-TBI datasets [28]. Moreover, the presence of mild TBI in our P-TBI model allows the disentanglement of focal-injury-related and systemic-injury-related inflammatory responses. This difference is clearly seen in the sensitivity to the lack of C3 and/or CD14: the cytokine induction in the site of mild TBI is resistant to the loss of C3 and/or CD14, indicating a substantial redundancy in the DAMP signaling, whereas the cytokine induction in the apparently non-injured brain remains sensitive, mainly to C3 loss. A similar redundancy of DAMP signaling may be preventing the loss of C3 and CD14 from affecting peripheral tissue damage and systemic inflammation.

Complement activation takes place within minutes after polytrauma [8] and the release of multiple biologically active proteolytic fragments from C3 and C5 drives a massive activation of innate immunity cells, such as neutrophils and macrophages [9], and the release of other inflammatory cytokines [31]. Complement activation is critical for secondary damage to kidney and lungs in polytrauma [32,33]. CD14, on the other hand, acts as a co-factor contributing to the signaling of multiple TLR, which recognize not only pathogens-associated molecules (PAMPs) but also DAMPs [4]. CD14 has been implicated in the response to polytrauma and to secondary injury to kidney, intestine and liver [5,34]. Our findings of abolished microglial reaction to polytrauma in mice lacking C3 and CD14 are aligned with previously reported beneficial effects in isolated TBI obtained in *C3^−/−^* animals or upon complement depletion [18,35] and are in line with the reduced microglial inflammation observed in *CD14^−/−^* animals [36]. However, the study of single- and double-KO animals reveals a larger dependency on C3 in the cortex and striatum but the dependency on either C3 or CD14 in the hippocampus. Interestingly, simultaneous loss of C3 and CD14 does not ameliorate the damage to any other organ considered (in particular lungs, liver, pancreas, and vasculature) in our polytrauma model with HS, although beneficial effects have been reported upon combined C5 and CD14 inhibition in a porcine polytrauma model [5]. Thus, both C3 and CD14 are dispensable for the systemic inflammatory response to polytrauma in present murine model, likely because of the redundant activation of inflammatory pathways by the massive release of DAMP but appear necessary to mediate the cerebral inflammatory response in these conditions.

The contrast of brain samples from WT animals and from single *C3^−/−^*or *CD14^−/−^* or from double *C3^−/−^CD14^−/−^*KO reveals that while *CCL2* and *TNF* induction in the cortex depends largely on CD14 and C3 respectively, both cytokines are mainly dependent on C3 in striatum. Actually, in striatum we also detected some anti-inflammatory role of CD14, since *TNF* and *CCL2* mRNA upregulation is abolished in *C3^−/−^* but is re-instated in *C3^−/−^CD14^−/−^*(Fig. 7I and 8E), indicating an opposite effect of the loss of C3 and CD14; these findings are in agreement with previously reported pro- and anti-inflammatory effects of CD14 loss in the microglia [36]. Our findings extend the previously reported role of C3 in driving striatal neuroinflammation upon *Staphylococcus* infection [37] as well as of its role in mediating microglial reactivity in a hypoperfusion model [38]. In contrast, the response to LPS, which requires CD14, has been previously shown to be different in the cortex and striatum [39], with substantially higher neurotoxicity observed in the striatum. More broadly, sensitivity to LPS has been shown to exhibit strong regional variations across the brain, with *TNF* induction resulting up to five times higher in the cortex than in subcortical structures [40]. Overall, the differential effects of C3 and/or CD14 loss are not fully explained by the expression patterns of these molecules, with C3 playing a more decisive role. Although the striatum expresses much more C3 than the cortex, C3 is critical in both structures, even if they display a similar expression of CD14. Thus, when C3 is available, it represents the master regulator of the microglial response to polytrauma, and CD14 may even exert some anti-inflammatory role.

The present work displays some unavoidable limitations. First, we considered a single, early timepoint for the tissue sampling; because of the high severity of the injury, criteria set in place to avoid unnecessary suffering of the animals did not allow for longer time courses. It is anticipated that the neuroimmune trajectory of the P-TBI may then evolve toward a phase of resolution, either complete, although in more severe P-TBI models, this is not observed (Rowe et al., 2024) or most likely to a permanent post P-TBI inflammatory state. Second, we focused our analysis on microglial cells, since they are highly sensitive to inflammatory stimuli and therefore constitute suitable readouts for interventions. Astrocytes, resident macrophages, pericytes and even neurons can contribute to the cytokine milieu (Janova et al., 2016). The discrepancy between microglia-selective cytokine expression and whole-tissue expression. e.g., the effect of the loss of both C3 and CD14 is detected at microglial level in ipsilateral cortex but not at whole-tissue level, points toward additional contributions to cytokine induction that are not C3 and/or CD14 sensitive and may not respond to therapeutic interventions on complement.

Taken together, our data support the acute onset of neuroinflammation in the context of P-TBI, similar to other systemic inflammatory response conditions (Chung et al., 2023). The profile of the neuroimmune activation is uniquely dependent on cerebral C3 and, to a lesser extent, on cerebral CD14, whereas for systemic inflammation in P-TBI in this model their role appears dispensable. Interventions on C3 and/or CD14 may therefore be titrated depending on the occurrence of a cerebral focal injury or diffuse encephalopathy.

## Methods

### Experimental animals and genotypes

Mouse polytrauma experiments have been approved by the animal oversight committee at Ulm University and by the corresponding authorities (Regierungspräsidium in Tübingen; Reg. 1480). Male mice aged 14-16 weeks of four different strains (wild-type: C57BL6; *C3^−/−^*: B6.129S4-C3tm1Crr/J; *CD14^−/−^*: B6.129S4-Cd14tm1Frm/J [41] and *C3^−/−^/CD14^−/−^*: B6.129S4-C3tm1Crr/ Cd14tm1Frm/J) were generated by Prof. Egil Lien (UMASS, Boston, MA) and provided by Prof. Tom Eirik Mollnes (University of Oslo, Norway), additional wild-type mice (C57BL6) were purchased for Janvier-Labs (France).

### Polytrauma model

The polytrauma and haemorrhagic shock (P-TBI) murine trauma model, consisting of blunt thorax trauma (TXT), traumatic brain injury (TBI), closed unilateral diaphyseal femoral fracture and a haemorrhagic shock (HS) was performed as previously reported [42,43]. Briefly, mice of each genotype were randomized either into the P-TBI group or sham surgery group, followed by buprenorphine administration (0.05 mg/kg, subcutaneously) and sevoflurane anaesthesia (3.0% in 97% O2). The bilateral blunt thorax trauma was performed using a well-defined blast wave. The traumatic brain injury was induced by closed head weight drop with a weight of 333 g falling from a height of 2 cm on the parietal bone. The closed diaphyseal femoral fracture was performed on the right leg by a weight dropping device of 50 g falling from a height of 120 cm. Directly after the polytrauma, the left femoral artery and the right jugular vein were catheterized. The HS was started 1 h after polytrauma induction, by withdrawing blood from the femoral artery until the mean arterial pressure (MAP) dropped to 30±5 mmHg, which was maintained for an additional 1 h. Following the haemorrhagic shock the mice received a 4-fold volume resuscitation of the withdrawn blood volume, with isotonic electrolyte solution (Jonosteril, Fresenius Kabi, Germany) over 30 min. After the fluid therapy, mice that could not reach a MAP above 50 mmHg, received norepinephrine (0.5-2.0 ug/kg/min). Control mice (sham group) underwent the same treatments (analgesia, anaesthesia, femoral artery and jugular vein catheters, and application of 400 ul of isotonic electrolyte solution) without the induction of the TXT, TBI, femoral fracture and HS. During the whole procedure animals were kept under general anaesthesia (sevoflurane), analgesia (Buprenorphine) and monitoring of heart rate (HR) and MAP using a blood pressure analyser (DSI, St. Paul, Minn).

### Blood and tissue sampling

Four hours after the induction of the P-TBI or sham surgery, mice were sacrificed followed by organ and blood harvesting for further processing. EDTA blood was drawn by cardiac puncture, followed by centrifugation for 5 min at 500 x g at 4°C and storage at −80°C. The brain was dissected and snap frozen on dry ice, brain tissue was either used for protein isolation for antibody arrays or RNA isolation for RT-qPCR.

A cohort of mice (only wild-type and *C3^−/−^/CD14^−/−^*) were anesthetized with a lethal dose of ketamine (100 mg/kg) and Xylazine (16 mg/kg) in saline followed by transcardial perfusion with 25 mL of PBS and 50 mL of 4% PFA. Brain tissue was dissected and postfixed overnight at 4°C in 4% PFA followed by washing in PBS and cryoprotection in 30% sucrose for 2 days. Brain samples were embedded in OCT (Tissue Tek) and stored at −80°C.

### Protein extraction and Antibody array analysis and bioinformatics

Proteins were isolated from the brain by adding RIPA buffer (150 mM NaCl, 50 mM Tris pH 7.6, 0.1% SDS, 1% Triton X-100, 1 mM EDTA), in a volume 4x the weight of the brain (weight: volume), protease and phosphatase inhibitors (Roche tablets) were added in excess, followed by homogenization of the tissue using a tissue grinder. The homogenate was sonicated for 2 x 2 sec at 65%, followed by incubation for 30 min at 4°C. The homogenate was centrifuged for 20 min at 10,000 x g, the supernatant was transferred to another tube and the pellet was discarded. The protein concentration was determined using the BCA assay (Thermo Fischer) according to the manufacturer’s instructions. Briefly, the homogenate was diluted 20x in RIPA buffer and added to the wells together with the standard in duplicates, 200ul of working reagent (50:1, reagent A:B) was added to each well and incubated for 30 min at 37°C. The absorbance was measured at 562 nm with a plate reader. Sample concentration was determined by calculation from the standard curve.

Targeted cytokine proteomics was performed using the mouse inflammation array G1 from raybiotech (Raybiotech), according to manufacturers’ instructions. Briefly, glass slides were removed from −20°C and equilibrated for 30 min at RT. Slides were removed from the package and air-dried for another 2 h, followed by blocking by adding 100 ul sample diluent to each well for 30 min at RT. Buffer was decanted and 100 ul of sample (400 ug) was added to the wells and incubated overnight at 4°C. Samples were decanted from the well and washed with 150 ul of 1x wash buffer I (diluted in dH2O) for 5 x 5 min at RT with gentle shaking, followed by another wash step with 150 ul of 1x wash buffer II (diluted in dH2O) for 2 x 5 min at RT with gentle shaking. Then, 80 ul of detection antibody cocktail was added to each well and incubated for 2 h at RT, followed by 5 x 5 min washing with wash buffer I and 2 x 5 min washing with wash buffer II. Then, 80 ul of Cy5 streptavidin (1:1000 in sample diluent) was added and incubated for 1 h at RT, followed by a 5 x 5 min washing with wash buffer I. The holders were disassembled and the glass slides were washed once for 15 min in an excess of wash buffer I (30 mL) and 5 min in an excess of wash buffer II (30 mL). After drying using a centrifuge at 1000 rpm for 5 min, the slides were scanned using a GenePix 4000B array scanner (Molecular Devices, LLC).

Antibody array analysis was performed using the R-studio version 4.3.1, using the limma package, utilizing linear models to model for experimental designs. Normalization within and between arrays was done using the cyclicloess method, to remove intensity-dependent biases between different samples and thereby making them more comparable. Then the arrays were corrected for any batch effects using the ComBat function, which estimates the mean and variance for each batch and adjusts the data accordingly. Then we proceeded towards differential expression, employing empirical Bayes methods that samples information across all proteins on the array, moderating the standard errors and improving the power to detect differential expression. The output was obtained using the topTable function to extract lists of differentially expressed proteins and their associated statistics like log fold-change, p-values, etc. as in Supplementary file 1.

### ELISA assay

The Enzyme-Linked Immunosorbent Assays (ELISA) were performed according to manufacturer’s instructions, with following dilutions, for Syndecan 1:10, I-FABP 1:100, GFAP 1:2 (LSbio), PAI-1 1:50, C5a 1:100 (R&D systems), LIPASE 1:5, C3b 1:1000 (Hycult), IL-6 1:2 (BD Bioscience), ALT 1:100 (Abcam), CC16 1:500 (Abbexa).

### RNA extraction and qPCR

Brain tissue was disrupted and homogenized in 1 ml of QIAzol (Qiagen, Germany). Following this, 200 µl of chloroform was added, and the mixture was vigorously vortexed for 15 seconds. After a 10-minute incubation at room temperature for phase separation, the homogenate was centrifuged at 12,000 × g for 10 minutes at 4 °C. The aqueous phase containing RNA was then transferred to a new tube. To precipitate the RNA, an equal volume of isopropanol was added to the aqueous phase, followed by a 10-minute incubation at room temperature. The mixture was centrifuged again at 12,000 × g for 10 minutes at 4 °C. The resulting RNA pellet was washed with 1 ml of 75% ethanol in DEPC-treated water and centrifuged at 8,000 × g for 10 minutes at 4 °C. The ethanol was carefully removed, and the RNA pellet was allowed to air-dry completely before being reconstituted in 30 µl of RNase-free water. RNA concentration and purity were quantified using a NanoDrop spectrophotometer, and the RNA was stored at −80 °C.

For reverse transcriP-TBIon, 1 µg of RNA was diluted to 40 µl with RNase-free water and 5 µl of random hexamers (Biomers, Germany) were added. After a 10-minute incubation at 70°C for primer annealing, the mixture was cooled on ice. A master mix containing reverse transcriptase, RNase Inhibitor, dNTPs mix, and M-MLV RT 5x buffer was added to the RNA-primer solution. The reaction mixture was incubated at room temperature for 10 minutes, then at 42°C for 45 minutes for cDNA synthesis. The reaction was terminated by heating at 99°C for 3 minutes, then immediately placed on ice. The resulting cDNA was stored at - 20°C.

Primers were designed using the Primer-BLAST tool from NCBI. Sequences from the NCBI GenBank database were input, and parameters were set for oP-TBImal PCR product size and primer melting temperature. Primers were synthesized by Biomers (Germany) and validated for specificity and functionality using test runs and controls. The detailed list of the primer sequences for each gene tested is reported in Supplementary Table 1. Quantitative RT-PCR (qRT-PCR) was conducted using the LightCycler 480 II system (Roche) with Power PCR TB Green PCR Master Mix (Takara, Japan). Each reaction included 2 µl of cDNA, 3 µl of primer mix, and 5 µl of TB Green in a 96-well plate. Samples were run in duplicate with GAPDH as an internal control. The threshold cycle value (Ct) was calculated and normalized using the equation 2 − ΔCt (ΔCt = Cttarget gene − CtGAPDH) to determine relative mRNA expression levels. Fold change was determined by comparing the experimental groups to the WT Sham group.

### Tissue sectioning and immunofluorescence staining

Brain tissue embedded in OCT were cryosectioned at 40 um using a cryostat, followed by blocking of the tissue in blocking buffer (3% BSA, 0.3% triton-X100, 1× PBS) for 2 h at room temperature (RT). Sections were incubated for 48 h at 4°C with primary antibodies (suppl. table 1) diluted in blocking buffer, followed by 3 x 30 min washing in PBS at RT. Sections were incubated for 2 h at RT with secondary antibodies (Supplementary Table 1) diluted in blocking buffer, followed by 3 x 30 min washing in PBS at RT. Finally, sections were mounted with Fluorogold prolong antifade mounting medium (invitrogen).

### Single mRNA in situ hybridisation

mRNA *in situ* hybridisation was performed according to manufacturer’s instructions (ACDBio, RNAscope, Newark, CA); fluorescent in-situ hybridization for fixed frozen tissue, all reagents and buffers were provided by ACDBio [44]. Briefly, 40 um tissue sections were mounted on superfrost plus glass slides-The slides were stored for at least 24h at −80°C and washed 2 x 5 min in 1 x PBS. The slides were baked for 30 min at 60°C, followed by a post-fixation in 4% PFA for 15 min at 4°C. The sections were dehydrated in a series of ascending ethanol concentrations (50% - 70% - 100%) for 5 min each, followed by a last time in 100% ethanol for 5 min. Sections were air dried and covered in 30% hydrogen peroxide for 10 min at RT, followed by 2x wash in PBS by moving the sections up and down 5x. Sections were incubated for 5 min in 1 x target retrieval solution at 100°C, followed by a short wash step in dH20 and twice in 100% ethanol. Sections were covered in protease III solution and incubated for 30 min at 40°C, followed by 2 x 2 min washing in dH20. The probes (C3, CD14, TNF and CCL2) were added and incubated for 2 h at 40°C, followed by a 2 x 2 min washing in wash buffer. Then, amplification 1 was added to the sections and incubated at 40°C for 30 min followed by 2 × 2 min washing step with wash buffer. Next, amplification 2 was added to the sections and incubated at 40°C for 30 min followed by 2 × 2 min washing step with wash buffer. As a final amplification step, amplification 3 was added and incubated for 15 min at 40°C, followed by 2 × 2 min washing step with wash buffer. HRP corresponding to the channel was added to the sections and incubated for 15 min at 40°C, followed by 2 x 2 min washing step with wash buffer. The diluted fluorophore (1:500 for all probes; in TSA buffer) was added to the sections and incubated for 30 min at 40°C, followed by 2 x 2 min washing step with wash buffer. The remaining HRP was blocked by adding HRP-blocker to the sections and incubated for 15 min at 40°C, followed by 2 x 2 min washing step with wash buffer. Sections were blocked using blocking buffer (3% BSA, 0.3% Triton X-100, PBS) for 1 h at RT, followed by primary antibody, diluted in blocking buffer, incubation overnight at 40°C. The sections were washed 3 x with PBS-T (1% triton-x100) at RT for 20 min each, followed by secondary antibody incubation (diluted in blocking buffer) for 2h at RT. The sections were washed 3 x with PBS-T at RT for 30 min each and mounted using Fluorogold prolong antifade mounting medium (Invitrogen).

### Image acquisition and analysis

Confocal images were acquired with a laser-scanning confocal microscope (LSM Leica DMi8 and Olympus Fluoview FV4000), with a 20x air objective (NA 0.75) and 40x oil objective (NA 1.3) in a 1024 × 1024-pixel resolution and a 12-bit (Leica) and 16-bit (Olympus) format. For the mRNA RNAscope imaging, a z-stack of 14-15 optical sections spanning 28-30 um (step size of 2 um), with a tilescan of 2 x 2 (40x objective) was made. For the microglia/pS6 imaging, a z-stack of 14-15 oP-TBIcal sections spanning 28-30 um (step size of 2 um), with a single tile (20x objective) was made. Imaging was done to obtain both the ipsilateral and contralateral side of the brain with 3 main regions (cortex, hippocampus and striatum). Imaging parameters were set to obtain signals from the stained antibody/probe while avoiding saturation. All fluorescent channels were acquired independently to avoid fluorescent cross-bleed.

Image analysis was performed using the ImageJ software. Images were background subtracted and smoothed. For the mRNA RNAscope analysis, images were contrasted to visualize all spots in the images, all spots per image were counted using the trackmate plugin (average spot size of 1 um). Additionally, TNF-⍺/CCL2 positive microglia were assessed manually with the cell counter plugin, positivity of microglia was determined by having at least 2-3 spots of TNF-⍺/CCL2 inside the soma of the microglia. For the immunofluorescence analysis, pS6 intensity per microglia was assessed by thresholding the microglia and using the plugin ‘analyze particles’ with saving all region of interests (ROIs). The ROIs were overlaid in the pS6 channel and intensity was assessed per ROI. The Scholl analysis was performed on single microglia with a start radius of 0.622 um, a step size of 1.864 um and end radius of 45 um (size of insert used), the number of intersections per step size were automatically assessed.

### Statistical methods

Data analysis was performed using the Graphpad prism version 8 software or R software suits Version 4.3.1. Statistical analysis was performed using animals as a biological unit; whenever appropriate, single data points and averages per animal are depicted in the graphs. All datasets were tested for normality using the Shapiro-Wilks test. When comparing two groups, an unpaired t-test or Mann-Whitney test was used. When comparing multiple groups, ANOVA with Sidak’s multiple correction or Kruskal-Wallis test with Dunn’s multiple correction were used, depending on normality. The comparison between the sham and P-TBI groups has been performed within each genotype, with analyses between genotypes when appropriate. Outliers were defined by the ROUT outlier test with a Q value of 1%. Significance was set at p < 0.05. Exact p values were reported where appropriate (with exception of the Scholl and hemodynamic analysis), descriptive statistical analysis with exact p values for all datasets are reported in supplementary information descriptive statistics.

## Conflict of interest

The authors declare no competing interests.

## Acknowledgement

We all thank the members of the CRC 1149 for their scientific input and discussion. We would like to thank Prof. Anita Ignatius for the access to the histology facility. We would like to thank Annette Palmer, Sonja Braumüller and Lena Dörfer for the mouse surgeries. Technical support by Thomas Lenk was highly appreciated.

## Funding

FR and MHL are supported by the Deutsche Forschungsgemeinschaft (DFG; Grant no. 251293561) in the context of the SFB1149/3 “Danger Response, Disturbance Factors and Regenerative Potential after Acute Trauma”. FR is also supported by DZNE core funding. FoH is also supported by the ZNS Hannelore Kohl Stiftung.

## Supplementary Figures

**Suppl. Figure 1: C3 and CD14 mediate diffuse, but not focal, P-TBI induced *TNF* and *CCL2* expression. A.** Experimental design of polytrauma and TBI in wildtype and *C3^−/−^/CD14^−/−^*mice, investigating bilateral cortex, hippocampus and striatum. **B-C.** RT-PCR analysis of *C3* and *CD14* expression in cortical tissue confirmed C3 and CD14 knockout mice. *CD14* expression revealed a significant increase in the WT sham vs WT P-TBI groups (WT sham vs WT P-TBI: p = 0.039). **D-G.** Single mRNA *in situ* hybridization of *TNF* revealed significant increase of *TNF* puncta in bilateral cortex (WT sham vs WT P-TBI; ipsi: p = 0.007; contra: p = 0.017), hippocampus (WT sham vs WT P-TBI; ipsi: p = 0.001; contra: p = 0.0099) and striatum (WT sham vs WT P-TBI; ipsi: p = 0.007; contra: p = 0.01) of WT mice. For *C3^−/−^/CD14^−/−^* mice, a significant increase of *TNF* puncta was observed in the ipsilateral cortex, hippocampus and striatum (*C3^−/−^/CD14^−/−^* sham vs *C3^−/−^/CD14^−/−^* P-TBI; cortex: p = 0.045; hippocampus: p = 0.048; striatum: p = 0.037), with no difference in the contralateral brain. **H-K.** Single mRNA *in situ* hybridization of *CCL2* revealed significant increase of *CCL2* puncta in bilateral cortex (WT sham vs WT P-TBI; ipsi: p = 0.003; contra: p = 0.016), hippocampus (WT sham vs WT P-TBI; ipsi: p = 0.005; contra: p = 0.037) and striatum (WT sham vs WT P-TBI; ipsi: p = 0.015; contra: p = 0.013) of WT mice. For *C3^−/−^/CD14^−/−^*mice, a significant increase of *CCL2* puncta was observed in the ipsilateral cortex, hippocampus and striatum (*C3^−/−^/CD14^−/−^*sham vs *C3^−/−^/CD14^−/−^* P-TBI; cortex: p = 0.01; hippocampus: p = 0.0002; striatum: p = 0.001), with no difference in the contralateral brain. Data shown as dot plots with all data points depicted and single mice as coloured dots (cyan; green). Scale bar: 200 um. Group size: WT sham N = 4, WT P-TBI N = 4. C3^−/−^/CD14^−/−^ Sham N = 3, C3^−/−^/CD14^−/−^mice P-TBI N = 4. \**p* < 0.05; \*\**p* < 0.01; ***p< 0.001; ns = not significant.

**Suppl. Figure 2: Loss of C3 and/or CD14 does not affect haemodynamic or systemic organ damage post P-TBI. A.** Experimental design of polytrauma and TBI (P-TBI) in wildtype, *C3^−/−^*, *CD14^−/−^*and *C3^−/−^/CD14^−/−^* mice, investigating both haemodynamic parameters (heart rate and mean arterial pressure) and plasma damage markers. **B.** Enzyme-linked immunosorbent assay (ELISA) measurement of C3b in plasma of all genotypes subjected to P-TBI confirmed C3 knockout. No difference was observed in P-TBI group. **C-D.** Mean arterial pressure (MAP) measurements in WT sham vs WT P-TBI revealed a drop in MAP upon HS initiation, which was kept stable for 1 h, followed by isotonic fluid therapy. No differences were observed between the genotypes. **F-H.** Heart rate measurements in WT sham vs WT P-TBI revealed a slight significant increase in heart rate around 180 min into the procedure. *C3^−/−^*, *CD14^−/−^*and *C3^−/−^/CD14^−/−^* revealed no overall difference in heart rate, with a small significant increase in the *CD14^−/−^* mice (compared to WT) at around 180 min into the procedure for both the sham and P-TBI groups. **I-K.** ELISA analysis of organ specific damage markers GFAP (brain), CC16 (Lung) and ALT (liver) in plasma post sham and P-TBI revealed significant increase in all genotypes (Sham vs P-TBI: GFAP; WT: p = 0.024; *C3^−/−^*: p < 0.0001; *CD14^−/−^*: p = 0.0004; *C3^−/−^/CD14^−/−^*: p = 0.0004, CC16; WT: p = 0.0002; *C3^−/−^*: p = 0.0007; *CD14^−/^*^−^: p = 0.0008; *C3^−/−^/CD14^−/−^*: p = 0.0001, ALT; WT: p = 0.0007; *C3^−/−^*: p = 0.002; *CD14^−/−^*: p < 0.0001; *C3^−/−^/CD14^−/−^*: p = 0.001). **L-N.** ELISA analysis of organ specific damage markers LIPASE (pancreas), I-FABP (Intestine) and syndecan (Vasculature) in plasma post sham and P-TBI revealed significant increase in all genotypes for LIPASE (Sham vs P-TBI; WT: p = 0.002; *C3^−/−^*: p = 0.003; *CD14^−/−^*: p = 0.002; *C3^−/−^/CD14^−/−^*: p = 0.017). For I-FABP a significant increase was found in WT and *C3^−/−^*, with trend in *CD14^−/−^* (Sham vs P-TBI; WT: p = 0.01; *C3^−/−^*: p = 0.033; *CD14^−/−^*: p = 0.098). For syndecan a significant increase was found in WT, *CD14^−/−^* and *C3^−/−^/CD14^−/−^*(Sham vs P-TBI; WT: p = 0.011; *CD14^−/^*^−^: p = 0.026; *C3^−/−^/CD14^−/−^*: p = 0.02). **O-Q.** ELISA analysis of systemic damage markers PAI-1, IL-6 and C5a in plasma post sham and P-TBI revealed significant increase in all genotypes for PAI-1 and IL-6 (Sham vs P-TBI: PAI-1; WT: p = 0.003; *C3^−/−^*: p = 0.001; *CD14^−/−^*: p < 0.0001; *C3^−/−^/CD14^−/−^*: p = 0.001; IL-6: WT; p = 0.008; *C3^−/−^*: p = 0.006; *CD14^−/−^*: p = 0.0003; *C3^−/−^/CD14^−/−^* : p = 0.048). For C5a a significant increase was found in WT and *CD14^−/−^* (Sham vs P-TBI; WT: p = 0.026; *CD14^−/^*^−^: p = 0.0019). Data shown as X-Y plots with mean and SD for each timepoint as well as bar plots with individual data points. Group size: WT sham N = 8, WT P-TBI N = 8, C3^−/−^ sham N = 8, C3^−/−^ P-TBI N = 9, CD14^−/−^ sham N = 7, CD14^−/−^ P-TBI N = 8, C3^−/−^/CD14^−/−^ sham N = 8, C3^−/−^/CD14^−/−^ P-TBI N = 8. \**p* < 0.05; \*\**p* < 0.01; ***p< 0.001; ****p< 0.0001.

**Supplementary Table 1: List of antibodies and RT-PCR sequences**

## Notes

### Competing Interest Statement

The authors have declared no competing interest.

