## Supplementary figures and images for "Loss of C3 and CD14 reduces region-specific neuroinflammation in a murine polytrauma model"

### Suppl. Figure 1

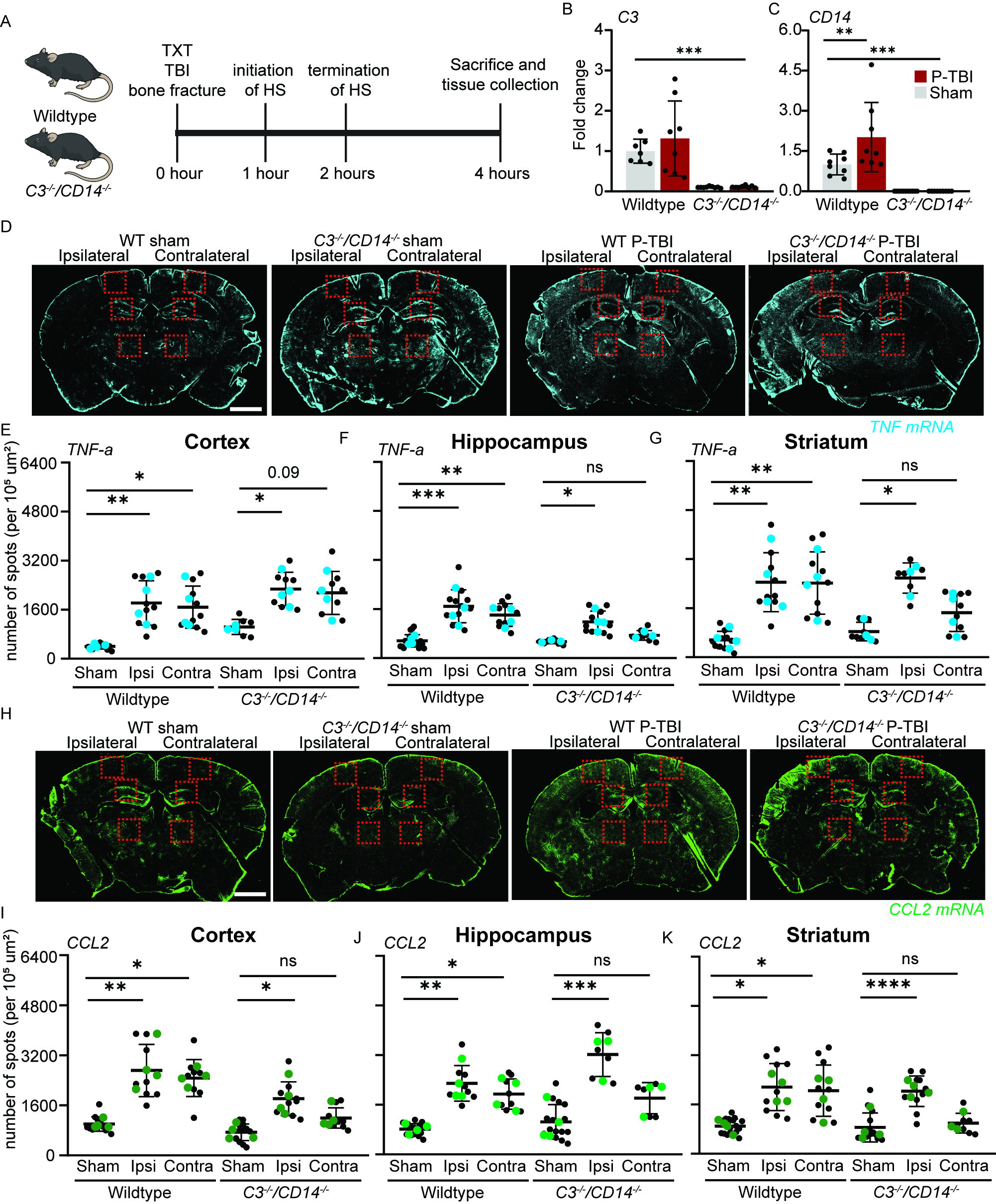

### Suppl. Figure 2

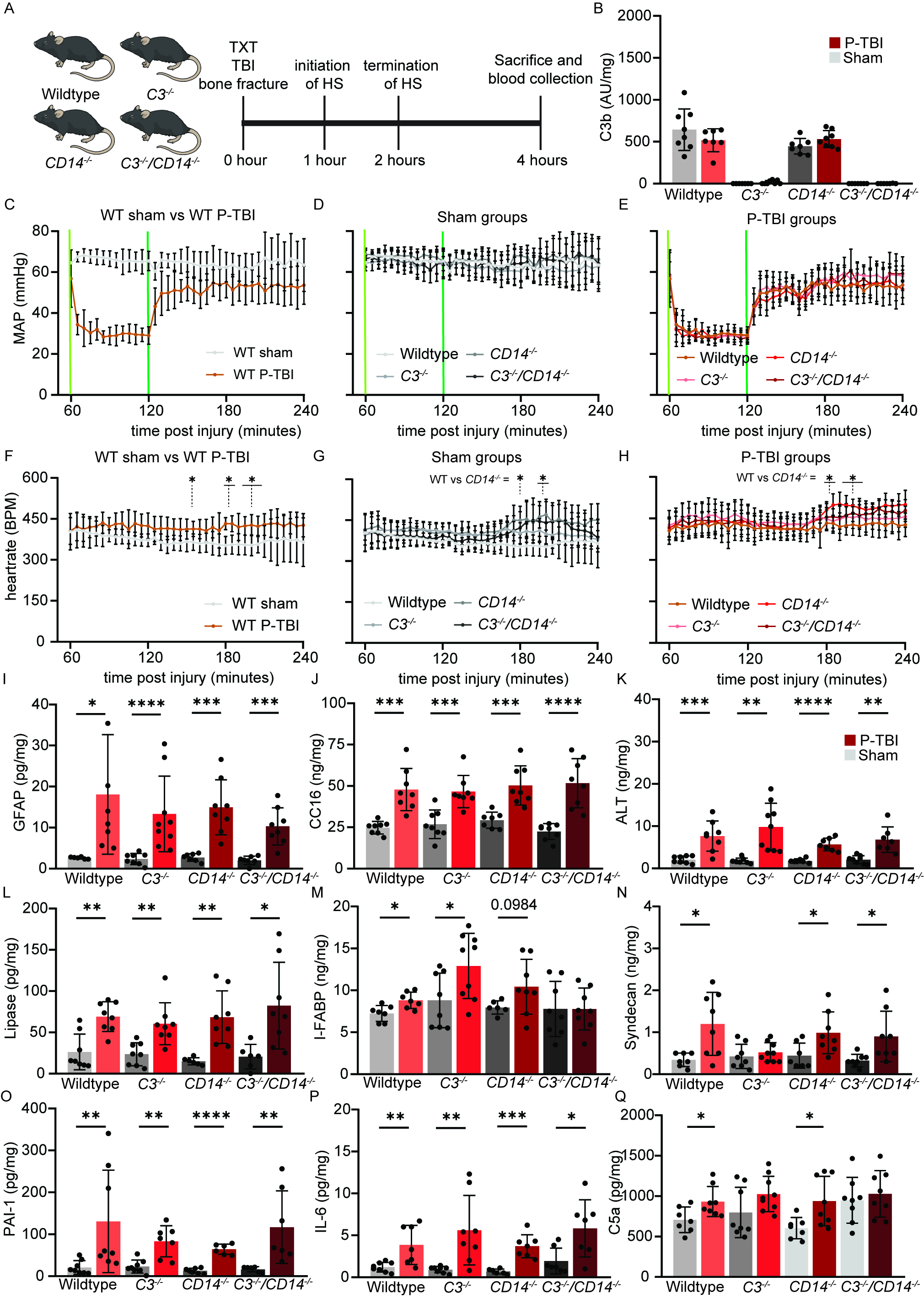
