## Supplementary material for "Loss of C3 and CD14 reduces region-specific neuroinflammation in a murine polytrauma model": Suppl. Table 1

**Supplementary Table 1: List of antibodies and RT-PCR sequences**

| **Antigen (primary antibody)** | **Catalogue number** | **Company** | **Dilution** |
| --- | --- | --- | --- |
| Rabbit anti IBA1 | 234008 | Synaptic Systems | 1:1000 (ISH) |
| Guinea pig anti IBA1 | 234004 | Synaptic Systems | 1:250 (IF) |
| Rabbit anti pS6-RP | 2211S | Cell Signaling Technologies | 1:200 (IF) |
| DAPI | D1306 | Thermo Fischer | 1:1000 (IF) |
| **Antigen (secondary antibody)** | **Catalogue number** | **Company** | **Dilution** |
| Donkey anti Rabbit AF 568 | A10042 | Invitrogen | 1:500 |
| Donkey anti Guinea pig CF 568 | 20377 | Biotium | 1:500 |
| Donkey anti Rabbit AF 647 | A21207 | Invitrogen | 1:500 |

| **Gene** | **Sequence** |
| --- | --- |
| TNF-a | Fwd: 5’ - TAG CCC ACG TGC AAC - 3’  Rev: 5’ - ACA AGG TAC AAT CGG C - 3’ |
| CCL24 | Fwd: 5’ – CAG CCT TCT AAA GGG GCC AA- 3’  Rev: 5’ – CTA AAC CTC GGT GCT ATT GCC- 3’ |
| CCL5 | Fwd: 5’ - TGC AGA GGA CTC TGA GAC AGC - 3’  Rev: 5’ - GAG TGG TGT CCG AGC CAT A - 3’ |
| CXCL13 | Fwd: 5’ – CTC CAG GCC ACG GTA TTC TG- 3’  Rev: 5’ – CCA GGG GGC GTA ACT TGA AT- 3’ |
| IL-7 | Fwd: 5’ – CTG CTG CAC ATT TGT GGC TT- 3’  Rev: 5’ – TGG CAA CTC TGT GAG ACT GG- 3’ |
| CCL2 | Fwd: 5’ - CCC AAT GAG TAG GCT GGA GA - 3’  Rev: 5’ - TCT GGA CCC ATT CCT TCT TG - 3’ |
| IL-1b | Fwd: 5’ - TGT AAT GAA AGA CGG CAC ACC - 3’  Rev: 5’ - TCT TC TTT GGG TAT TGC TTG G - 3’ |
| IL-33 | Fwd: 5’ - GGG CTC ACT GCA GGA AAG TA - 3’  Rev: 5’ - TTT GCC GGG GAA ATC TTG GA - 3’ |
| IL-6 | Fwd: 5’ - GCT ACC AAA CTG GAT ATA ATC AGG A - 3’  Rev: 5’ - CCA GGT AGC TAT GGT ACT CCA GAA - 3’ |
| CXCL1 | Fwd: 5’ – ACT CAA GAA TGG TCG CGA GG- 3’  Rev: 5’ – ACT TGG GGA CAC CTT TTA GCA- 3’ |
| CXCL2 | Fwd: 5’ – CAT CCA GAG CGT GAC G- 3’  Rev: 5’ – GGC TTCA GGG CAA ACT- 3’ |
| C3 | Fwd: 5’ – TCA CTA TGG GAC CAG CTT CAG- 3’  Rev: 5’ – AGC CGT AGG ACA TTG GGA GTA- 3’ |
| CD14 | Fwd: 5’ – CAT CTT GAA CCT CCG CAA CG- 3’  Rev: 5’ – TCG CAG GAA AAG TTG AGC GA- 3’ |
| GAPDH | Fwd: 5’ – AGC TTG TCA TCA ACG GGA AG- 3’  Rev: 5’ – TTT GAT GTT AGT GGG GTC TCG- 3’ |
